# *Chlamydia*-induced *ex vivo* activation of B cells from mice, nonhuman primates, and humans produces large numbers of uniformly activated antigen presenting cells

**DOI:** 10.1101/2025.07.25.666778

**Authors:** Rodolfo D. Vicetti Miguel, Joseph G. Charek, Shumin Fan, Dondrae Coble, Nirk E. Quispe Calla, Thomas L. Cherpes

## Abstract

*Chlamydia trachomatis* (*Ct*) is a clinically important pathogen that causes ocular and genital infections. While genital *Ct* infection in women is typically persistent and asymptomatic, the host responses that dampen inflammation while gradually eradicating this bacterium are only partially defined. Herein, we show that *Ct* promotes human B cell activation via TLR2-mediated signaling of MyD88-dependent pathways and non-antigen-specific B cell receptor signaling that engages the CD19 and CD79a/b complex. We also found that *ex vivo Chlamydia*-induced activation of B cells from the peripheral blood of humans or rhesus macaques (RM), or from murine splenocytes, generates cells with a phenotype characteristic of an activated antigen presenting cell (APC). Consistent with this phenotype, intravenous injection of *Chlamydia*-activated B cells (CAB) loaded with various cognate antigens (Ag) elicited robust Ag-specific T cell immunity in several experimental models. Specifically, RM injected with CAB loaded with a model Ag developed Ag-specific CD8^+^ T cell immunity and mice injected with Ag-loaded CAB formed CD8^+^ T cell immune responses that protected against virus challenge and tumor development. As *Chlamydia* nonspecifically activated B cells to form large numbers of uniformly activated APC readily loaded with cognate Ag, our studies suggest that CAB may offer foundation for a cellular vaccine platform that elicits robust Ag-specific CD8^+^ T cell immunity against a variety of tumors and microbial pathogens.

## INTRODUCTION

Dendritic cells (DC), macrophages, and B cells are professional antigen-presenting cells (APC) that coordinate the response of the innate and adaptive arms of host immunity (1). DC serve as sentinel cells and APC for naïve T lymphocytes, and there are longstanding efforts to develop DC-based vaccines that stimulate antigen (Ag)-specific immunity against tumors and microbial pathogens (1, 2). Clinical research shows that DC-based vaccines elicit T cell responses to tumor-associated Ag, but none to date significantly improved primary study outcomes, including overall rates of survival (3, 4). While checkpoint inhibitors and other immunotherapies are likely to enhance the efficacy of DC-based vaccines (5), it may also be important to remove or reduce limitations of the platform itself. These limitations include problems producing sufficient numbers of autologous DC for multiple treatments and creating methods to manufacture and administer these vaccines that are less labor and resource intensive than those currently available (6–11).

Compared to DC, B cells are a larger pool of peripheral blood APC more amendable to *ex vivo* expansion (12). While B cells were found more likely to induce tolerance than boost Ag-specific immunity (13, 14), these results were mainly acquired with resting or lipopolysaccharide (LPS)- activated B cells (15, 16). Of note, stimuli other than LPS elicit greater B cell proliferation and costimulatory molecule expression (12, 17, 18) and capacity of B cells to function as APC and promote Ag-specific T cell immunity is dependent on the strength and nature of B cell activation (19). Moreover, T cell-independent microbial antigens induce non-specific B cell activation via pathogen-associated molecular pattern signaling or binding of the B cell receptor (BCR) outside sites of Ag-binding (20). *Trypanosoma cruzi*, *Plasmodium falciparum*, and *Staphylococcus aureus* are among the pathogens that polyclonally activate B cells and promote nonspecific antibody production, responses that promote pathogen persistence during acute infection (20). While less explored, the intracellular bacterium *Chlamydia trachomatis* (*Ct*) may also use non-specific B cell activation as an immunoevasive response. *In vitro* studies show that *Ct* promotes B cell proliferation and polyclonal antibody production (21) and hyperglobulinemia is identified in neonates diagnosed with pneumonia caused by *Ct* (22). B cell- and plasma cell-rich lymphoid follicles are also seen in *Ct*-infected ocular and genital epithelium (23) and endometrial tissue in women genitally infected with *Ct* displays significantly increased B cell numbers and expression of genes linked to B cell development (24–26). Consistent with promoting a nonproductive host response, protective immunity in women with untreated genital *Ct* infection typically requires weeks or months to form (27–29). These observations suggest that *Ct* promotes non-specific B cell activation, and herein we sought to define *Ct*-mediated mechanisms of B cell activation and characterize *Ct*-activated B cells (CAB), including the expression of costimulatory and adhesion molecules. We also explored CAB-mediated priming of Ag-specific CD8^+^ T cell immunity and the use of Ag-loaded CAB to induce *in vivo* immune responses against tumor and microbial Ag.

## MATERIALS AND METHODS

### Mice, cell lines, and microorganisms

Mouse studies were reviewed and approved by the Ohio State University (OSU) Institutional Animal Care and Use Committee (IACUC) and were carried out in accordance with the Guide for the Care and Use of Laboratory Animals and relevant national and institutional regulations. All methods are reported in accordance with the ARRIVE guidelines. Where indicated, 6-10-week-old C57BL/6J (B6), BALB/cJ, B6.129P2(SJL)-Myd88^tm1.1Defr^/J (MyD88^-/-^) (30), C57BL/6-Tg(TcraTcrb)1100Mjb/J (OT-I), and B6.Cg-Thy1^a^/Cy Tg(TcraTcrb)8Rest/J (pmel-1) mice (Jackson Laboratories, Bar Harbor, ME) were housed in specific-pathogen-free conditions. B16-F10 (CRL-6475) and E.G7-OVA (CRL-2113) tumor cell lines were obtained from ATCC (Manassas, VA, USA) and maintained as described (31, 32). HSV-2 strain 333 was also maintained using described methods (33) and *Ct* lymphogranuloma venereum (LGV) serovar L2 (VR-902B), *Ct* serovar D (VR-885), and *C. muridarum* (VR-123) were purchased from ATCC and maintained as described earlier (34, 35). Inactivation of *Chlamydia* elementary bodies (EB) and inclusion forming units (IFU) calculations were performed as detailed elsewhere (24, 36). For all animal experiments, healthy animals without signs of pre-existing disease were included. Animals were excluded if they exhibited signs of illness unrelated to experimental procedures. Mice were housed in groups of 3-5 per cage in individually ventilated cages with standard bedding, nesting material, and enrichment items. Ambient temperature was maintained at 20-22°C with relative humidity of 50-60% and 12h/12h light/dark cycles. Mice were randomly allocated to experimental groups using a computer-generated randomization schedule and cage mates were distributed across different experimental groups to minimize cage effects. All procedures were performed during the light phase of the light/dark cycle. Mice were acclimatized to the housing facility for at least 7 days before experiments began. Prior to sample collection or experimental procedures, mice were acclimatized to the procedure room for at least 30 minutes. At study termination, mice were euthanized in accordance with American Veterinary Medical Association (AVMA)-approved guidelines and OSU IACUC-approved protocol using carbon dioxide asphyxiation followed by cervical dislocation to ensure death. Animal death was verified prior to tissue collection and disposal.

### B cell activation studies

Human buffy coats were obtained from the Central-Southeast Ohio Region American Red Cross as approved by their Institutional Review Board, in accordance with relevant guidelines and regulations, including the Declaration of Helsinki. Informed consent was obtained from all donors. Peripheral blood from rhesus macaque (RM) (*Macaca mulatta*) were obtained as described in RM CAB vaccination studies section below. Peripheral blood mononuclear cells (PBMC) or splenocytes were isolated from human buffy coats or MR whole blood and mouse spleens, respectively, and processed as detailed earlier (37, 38). Also as described (39, 40), PBMC and splenocytes were cultured in complete medium for 4 days at 37°C and 5% CO2 with inactivated EB at specified multiplicities of infection (MOI) that were based on IFU calculated from *Chlamydia* stock prior to inactivation. Of note, generation of CAB from peripheral blood of humans or RM was unaffected by prior or current *Ct* infection (Figures S1C and S1E). A MOI of *Ct* L2 stock that elicited EC50 responses was used to delineate the signaling pathways induced by *Chlamydia*-induced B cell activation. B cell proliferation was normalized by designating positive controls to have 100% proliferation. *In vitro* human Toll-like receptor (TLR) screening was performed using a panel of HEK293-TLR-Blue clones as previously detailed (41). The primary TLR ligand screening was completed by InvivoGen (San Diego, CA, USA). For studies with blocking antibodies or chemical inhibitors, human PBMC were pre-incubated for 30 minutes at 37°C with indicated molecules, including neutralizing IgA antibody against TLR2 (B4H2) and TLR5 (Q2G4) (InvivoGen). Antibodies against the surface portions of CD79a (JCB117, Santa Cruz Biotechnology), CD79b (CB3-1, eBioscience), and CD19 (HIB19, BioLegend, San Diego, CA) blocked BCR-complex associated molecules. ODN2006 (InvivoGen) was used to induce TLR9-dependent proliferation of human B cells. Stimulation with anti-human IgM+IgG+IgA F(ab’)2 fragments (α-BCR) or Pam3CSK4 (TLR2/1 agonist) was performed as described (42). Kinase inhibitors R406 and BIIB-057 (Cayman, Ann Arbor, MI, USA); Ly294002 and PD98059 (InvivoGen), ibrutinib (Selleckchem, Houston, TX, USA), and the MyD88 dimerization inhibitor ST2825 (ChemScene, Monmouth Junction, NJ, USA) were used to pre-treat human PBMC. To define the bacterium components that induced B cell activation, inactivated *Ct* L2 EB were incubated with select antibodies for 30 minutes at 37°C prior to incubation with human PBMC. These antibodies included mAb against *Ct* LPS, clone CL21-335.2.3. (IgG2a, Abcam, Cambridge, MA) and the following anti-*Ct* MOMP antibodies: polyclonal, goat polyclonal IgG against *Ct* L2 MOMP (Bio-Rad, Hercules, CA); mAb 1, clone HT10 (IgG2a, Abcam); and mAb 2, clone 1297/143 (IgG2a, Abcam). For Western blots (WB), human B cells were purified by negative immunomagnetic selection according to manufacturer’s instructions (STEMCELL Technologies, Tukwila, WA). Phospho-protein WB were performed as described previously, signal detection using X-ray film and quantification using ImageJ software applying only minimal and uniform adjustments across entire images and equally to experimental and control samples (43). No images from different times or locations were combined into single. Unprocessed full images Western blots are included in the Supplementary Information (Figure S5).

### T cell priming studies

For labeling studies, cells re-suspended at 2x10^7^/mL in PBS containing 0.2% FBS were mixed with PBS that contained CellTrace Violet (CTV, Invitrogen) (10 µM for murine cells and 5 µM for human cells and prepared using manufacturer’s instructions). Suspensions were incubated for 12 minutes at 37°C, mixed after adding 4 volumes of complete medium, and incubated at room temperate for another 3 minutes. Cells were washed by centrifugation and re-suspended in complete media or PBS for *in vitro* or *in vivo* use, respectively. Assays to define allogeneic naïve T cell stimulation were done as already detailed (39) and flow cytometric evaluation performed on days 4 and 7 of incubation for murine and human T cells, respectively. Human CABs generated as described above were in some instances loaded with a CEF peptide pool (Thinkpeptides ProImmune, Oxford, UK) and incubated with CTV-labeled autologous memory CD8^+^ T cells, which were enriched using the EasySep™ Human Memory CD8^+^ T Cell Enrichment Kit (STEMCELL Technologies). These co-cultures were incubated for 7 days and replenished with fresh medium on days 3 and 6. Proliferation of Ag-specific memory CD8^+^ T cells was assessed via CTV dilution, and effector function was evaluated after restimulation of the same cultures with CEF peptide pool in the presence of brefeldin A during the last 6 hours of incubation, followed by intracellular cytokine staining (ICS). To define murine T cell priming, CD8^+^ T cells from OT-I or pmel-1 mice were purified by negative immunomagnetic selection (STEMCELL Technologies) from CTV-labeled splenocytes and 10^6^ CD8^+^ T cells were intravenously (i.v.) administered to B6 mice. The next day, indicated numbers of resting B cells, unloaded CAB, CAB loaded overnight with 5 µg/mL of EndoFit ovalbumin (OVA, InvivoGen), or CAB loaded 2 hours with 1 µM of one of the following peptides: OVA257-264 (SIINFEKL) (InvivoGen) or gp10025-33 (KVPRNQDWL) (Thinkpeptides ProImmune, Oxford, UK) were i.v. transferred to mice in 200 µL of PBS. To compare efficacy of T cell priming in fresh vs. cryopreserved CAB, B6 mouse splenocytes were stimulated for 4 days with inactivated doses of *Ct* L2 EBs shown to produce maximum B cell proliferation (cultures were replenished with fresh media and EndoFit OVA on day 3 of incubation). For fresh CAB, after stimulation and loading, cells were washed with PBS and re-suspended at 5x10^7^ CABs/mL in PBS. For cryopreserved CAB, identically handled cells were cryopreserved after a 4-day stimulation, thawed, and washed to remove cryopreservation media immediately before use. Mice i.v. vaccinated with either CAB and spleens were collected 3 days later and processed into single-cell suspensions. Splenocytes were stained as detailed in methods below and flow cytometric evaluation of Ag-specific T cell proliferation performed immediately (i.e., without fixation).

### Flow cytometry

Surface and intracellular staining for phenotyping and ICS were performed as reported earlier (24, 34, 37). To discriminate live vs. dead cells, LIVE/DEAD™ Fixable Near-IR dead cell stain kit (Invitrogen) was used. Blocking of non-specific binding prior to surface mAb staining was performed using TruStain FcX™ (anti-mouse CD16/32) antibody, Human TruStain FcX™ blocking solution (both BioLegend) or Human BD Fc Block™ (BD Biosciences, San Diego, CA) for mouse, human or NHP cells, respectively in combination with purified Mouse IgG1 isotype antibody. In addition, human and NHP cells were also blocked with human IgG isotype control (Invitrogen). In studies using human cells, the following fluorochrome-conjugated mAb were used: CD16 (3G8), CD25 (M-A251), CD40 (5C3), CD80 (L307.4), CD86 (2331), IgG (G18-145) and HLA-ABC (G46-2.6) (BD Biosciences, San Diego, CA); CD2 (RPA-2.10), CD3ε (UCTH1), CD3ε (HIT3a), CD4 (SK3), CD4 (OKT4), CD14 (M5E2), CD19 (HIB19), CD20 (2H7), CD27 (0323), CD124 (G077F6), CD137L (5F4), CD252 (11C3.1), IgD (IA6-2) and IgM (MHM-88) (BioLegend); CD5 (L17F12), CD8α (SK1), CD14 (61D3), CD20 (L26), CD54 (RR1/1), CD58 (TS2/9), CD62L (DREG-56), CD79a (HM47), CD95 (DX2), CD138 (DL-101), IFN-γ (4S.B3), TNF (MAb11), HLA-DR (L243) (eBioscience, San Diego, CA); and CD66abce (TET2) (Miltenyi Biotec, Auburn, CA). Phosphoflow of human PBMC was performed as previously described (44) with the following mAb: CD3 (SK7), CD20 (H1) (BD Biosciences), pSyk Y348 (moch1ct), and pBtk Y551 (M4G3LN) (eBioscience). For studies with murine cells, the following fluorochrome-conjugated mAb were used: CD4 (RM4-5), CD4 (GK1.5), CD8α (53-6.7), CD25 (PC61.5), CD40 (1C10), CD45R (RA3-6B2), CD62L (MEL-14), CD80 (16-10A1), CD86 (GL1), CD90.1 (HI551), CD90.2 (30-H12), CD137L (TKS-1), CD252 (BM134L), MHC-II (M5/114.15.2), MHC-I (AF6-88.5.5.3), NK1.1 (OK136), FoxP3 (FJK-16S), Vα2 (B20.1) and Vβ13 (MR12-3) (eBioscience); CD11b (M1/70), CD11c (N418) and Gr-1 (RB6-8C5) (BioLegend); CD54 (3E2), Ly6G (1A8) and Ly6C (AL-21) (BD Biosciences). MDSC gating was performed using previously suggested recommendations (45). The following fluorochrome-conjugated mAb were used with RM cells: CD2 (RPA2.10, eBioscience), CD3ε (SP34-2, BD), CD4 (L200, BD), CD8α (RPA-T8, BioLegend), CD14 (M5E2, BD Biosciences), CD16 (3G8, BD Biosciences), CD19 (J3-119, Beckman Coulter), CD20 (L27, BD Biosciences), CD40 (5C3, BD Biosciences), CD79a (HM47, eBioscience), CD80 (L307.4, BD), CD86 (IT2.2, BioLegend), CD86 (2331, BD), HLA-DR (L243, BioLegend), HLA-DR (TU36, BD), IFN-γ (B27, BD), IL-4 (8D4-8, eBioscience) and TNF (Mab11, eBioscience). To define Ag uptake *in vitro*, murine CAB or resting B cells were isolated by immunomagnetic selection (STEMCELL Technologies), resuspended in complete medium, and incubated with indicated concentrations of OVA-AF647 (Invitrogen, Carlsbad, CA) for 1 hour at 4°C or 37°C. To evaluate *in vitro* Ag processing, murine CAB or resting B cells were incubated with 10 μg/ml of DQ-OVA (Invitrogen) for varying times at 4°C or 37°C. Cells were washed with PBS before adding 7-AAD for flow cytometric evaluation. As indicated, integrated median fluorescence intensity (iMFI) was calculated as described (38). Cells were collected using LSR-II and LSRFortessa™ cell analyzers (BD Biosciences) supported by the OSU Comprehensive Cancer Center (OSUCCC) Analytical Cytometry Shared Resource and data was analyzed using FlowJo software (BD Biosciences) (fluorescence minus one controls were used to establish gates and confirmed with isotype controls).

### RM CAB vaccination studies

RM studies were approved by the OSU IACUC performed in accordance with the Guide for the Care and Use of Laboratory Animals and relevant national and institutional regulations. All methods are reported in accordance with the ARRIVE guidelines. Three female adolescent Chinese rhesus macaques (RM) testing negative for Macacine herpesvirus 1, group D simian retrovirus (serotypes 1, 2, 3, 4 and 5), simian immunodeficiency virus, simian T-cell leukemia virus types 1 and 2, *Chlamydia felis*, and *Chlamydia psittaci* were purchased from Primgen (Hines, IL). Upon arrival to our facility, RM underwent physical examination and 60-day quarantine (that included tuberculosis screening). They were housed in accordance with the Guide for the Care and Use of Laboratory Animals, with a standardized diet and environmental enrichment including toys and visual stimulation, and were provided with 24-hour rest periods between sampling procedures to minimize stress. At the time of study initiation, RM were between 4.5 - 5.5 years of age and weighed between 7.8 – 9.0 kg. Serum aliquots were used to confirm absence of circulating antibodies (IgG) against *Ct* serovar D (VR-885) and *Ct* LGV serovar L2 (VR-902B) (both ATCC) via in-house indirect whole cell inclusion immunofluorescence assay (24, 36). For indicated procedures, RM were anesthetized by intramuscular injection of ketamine (10 mg/kg). Prior to the administration of CAB, RM had been genitally infected with *Ct* serovar D for an unrelated study. Of note, we performed studies in mice that confirmed the presence of *Chlamydia*-specific immunity did not affect the induction of CD8^+^ T cell responses by Ag-loaded CAB. (Figure S1F). Using the methodology detailed above, NHP autologous CAB were generated and negative immunomagnetic selection performed using PE-conjugated mAbs against CD3ε (SP34-2), CD14(M5E2) and CD16(3G8) (BD Biosciences, San Diego, CA) and EasySep™ Other Species PE Positive Selection Kits from STEMCELL Technologies (Figure S4G-H). Anti-CD2 antibody (clone RPA-2.10) was excluded from this antibody pool for immunomagnetic selection because of its binding to human and RM CAB (Figure S4A-F). Immediately prior to the infusion of CAB, diphenhydramine (0.9 mg/kg) was i.v. injected and RM monitored for symptoms of distress and temperature fluctuations (no significant changes occurred). At study termination, RM were euthanized by intravenous administration of an overdose of sodium pentobarbital, consistent with AVMA guidelines for the euthanasia of laboratory animals and our OSU IACUC-approved protocol. After death was verified, spleens were collected and processed into single-cell suspension as previously described (38).

### Mouse CAB vaccination studies

For prophylactic immunization, B6 mice received 3 i.v. doses of 10^7^ CABs loaded with indicated protein (OVA) or immunodominant peptides (gB498-505, SSIEFARL; TRP-2180-188, SVYDFFVWL; gp10025-33, KVPRNQDWL; all Thinkpeptides ProImmune) at days 0, 5, and 7. HSV-2 infection and tumor challenge were performed 9 days after initial CAB infusion. For HSV-2 infection, corneas of anesthetized mice were scarified 6 times with a 30G needle before inoculation with 3x10^3^ PFU of HSV-2 333 (33). *In vivo* tumor experiments using B16-F10 and EG.7-OVA tumor cells were performed as already described (31, 32). For therapeutic immunization, B6 mice were subcutaneously (s.c.) injected with 7.5 x 10^5^ E.G7-OVA tumor cells. When tumors were palpable (i.e. day 5 after injection), mice were administered a total of 5 doses (injected every other day) of 10^7^ OVA-loaded CAB. Morbidity and tumor size (32) were monitored daily and tumor growth compared by area under the curve (AUC) analyses. In other studies, mice were euthanized 18 days after E.G7-OVA tumor inoculation, and tumors and spleens collected for tumor weight measurements and flow cytometric analyses of myeloid-derived suppressor cells (MDSC) and regulatory T cell populations, respectively. where indicated in these studies, CD8^+^ T cells were depleted using mAb as described previously (35).

### Statistical considerations

Investigators were blinded to group allocation during the experiment and outcome assessment wherever feasible. Sample sizes for mouse experiments were determined based on power calculations using preliminary data for each primary endpoint and performed using G*Power 3.1.9.6 software (46, 47). For RM experiments, the sample size (n=3) was chosen based on feasibility constraints while still allowing for statistical comparisons. In all *in vivo* experiments, the individual animal (mouse or rhesus macaque) was considered the experimental unit. For *in vitro* experiments using primary cells, individual cell cultures derived from separate animals were the experimental units. All experiments included appropriate control groups as detailed for each specific study. No animals or data points were excluded from the analyses unless pre-specified criteria (illness unrelated to experimental procedures, technical failure) were met and all animals/data points were included in the analyses unless otherwise specified in the figure legends. Where adequate, primary/secondary outcomes are outlined in the results section. All statistical analyses were performed using Prism 10 software (GraphPad, La Jolla, CA). Data normality was tested, depending on sample size, by the D’Agostino–Pearson omnibus K2 test, the Shapiro-Wilk test, or evaluation of the residuals. Differences between 2 independent groups were compared by unpaired Student’s t test. For comparison between paired samples, paired Student’s t tests were used. For assessment of tumor growth, AUC for individual tumor growth was calculated (tumor size [mm^2^] x days after tumor cell injection) and AUC values compared. For multiple group comparisons, depending on data distribution, one-way ANOVA and Tukey’s or Dunnett’s multiple comparison post hoc tests were used. *P* values < 0.05 were considered statistically significant and denoted in figures to highlight important between-group differences.

## RESULTS

### *Chlamydia* nonspecifically activates B cells

While reports of *Ct*-induced stimulation of human and mouse B cell proliferation and polyclonal immunoglobulin secretion were first published 40 years ago (48, 49), mechanisms underlying this effect, magnitude of the response, and phenotypic characterization of *Chlamydia*-activated B cells (CAB) was unexplored. We began the current study by exploring response magnitude, incubating human PBMC with variable amounts of inactivated *Ct* L2 EB. These studies identified that *Ct* L2 elicits robust and dose-dependent B cell proliferation (Figures 1A & S1A) and that similar levels of B cell proliferation were seen when human PBMC were incubated with EB from inactivated *Ct* serovar D or live *Ct* L2 (Figure S1B). Initial studies also suggested that *Ct*-induced proliferation did not require B cell infection, as no viable *Ct* was detected after CAB generation (Figure S1G). This result is also consistent with prior report that *Ct* does not productively infect B cells or T cells (50). Initial *ex vivo* studies also showed *Ct-*induced human B cell proliferation is not dependent on *Ct*-specific T cells (Figure S1C) and that B cell proliferation is associated with increased cell size and granularity (Figure S1A) and increased surface expression of IgG, CD5, CD95, CD124, and CD19/CD20 (Figure 1B & S1D). Compared to resting B cells, human CAB also displayed increased expression of CD40, CD80, CD86, CD137L (4-1BBL) and CD252 (OX40L), MHC molecules HLA-DR and HLA-ABC, and the adhesion molecules CD54 (ICAM-1), CD58 (LFA-3) and CD62L (L-selectin) (Figure 1C & 1D). Moreover, CAB demonstrated uniform upregulation of these costimulatory and adhesion molecules (Figure 1C-E).

**Figure 1.**
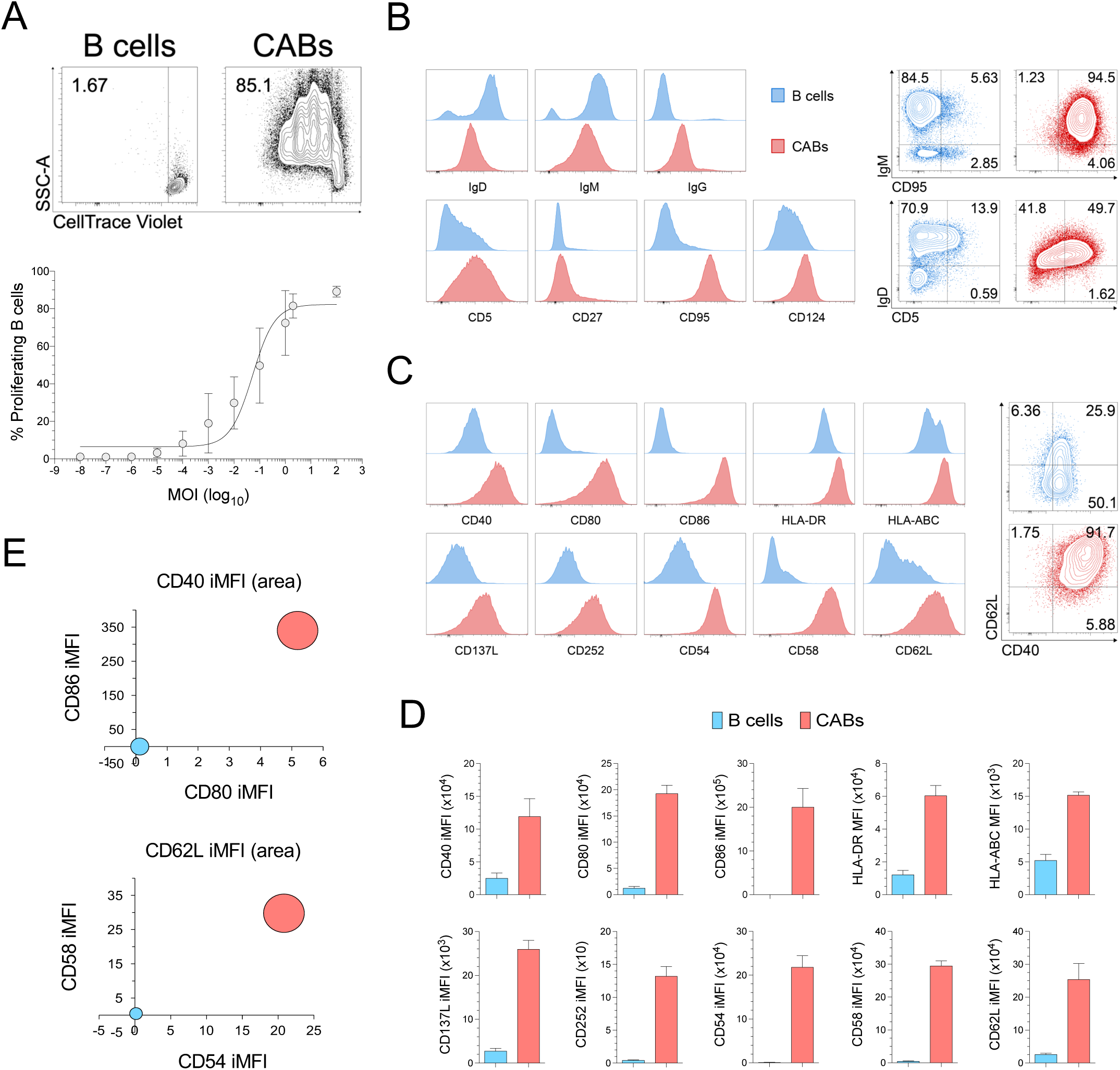
*Ct* nonspecifically activates human B cells. A) Fluorescently-labeled human PBMC were incubated for 4 days with varying amounts of inactivated *Ct* L2 elementary bodies to define proliferation of B cells (defined as live CD3^neg^CD14^neg^CD19^+^CD20^+^ cells) by flow cytometry. The top panel depicts representative contour plots of fluorescent tracer dilution in B cells incubated with vehicle (B cells) or *Ct* (CAB). Lower panel depicts dose-responsive B cell proliferation; MOI (multiplicity of infection) (mean ±SD). B & C) Representative histograms and contour plots show expression of (B) B cell markers and (C) adhesion and costimulatory molecules by B cells (blue) vs. CAB (red) (quadrant numbers denote percentages). D) Relative B cell vs. CAB expression for molecules shown in (C); bars denote mean ± SD. E) Comparing the relative expression of CD40, CD80 and CD86 (top panel) and CD54, CD58 and CD62L (lower panel) in B cells (blue) vs. CAB (red), bubble charts illustrate the uniform CAB-mediated induction of these molecules (iMFI expressed as x10^4^ for all markers). Data was acquired from 3-10 separate experiments, with between-group differences in panel D performed using unpaired Student’s tests.

Similar to the effect in human B cells, *ex vivo* stimulation of mouse splenocytes with inactivated *Ct* EB (serovar L2 or D) or inactivated *C. muridarum* (*Cm*) EB promoted dose-dependent B cell activation (Figures 2A & S2A-D), and murine and human CAB displayed comparably increased levels of MHC and costimulatory molecule expression (Figure 2B & 2C). 4 days after initiating *Ct*-induced stimulation, mouse splenocyte cultures consisted entirely of CAB and compared to resting B cells, CAB had higher levels of CD45R, MHC-II and CD54 levels and lower levels of CD62L (Figures 2B & 2C; S2B-S2E). Incubation of PBMC from RM with variable amounts of inactivated *Ct* L2 EB likewise indicated that *Ct* elicits nonspecific B cell proliferation (Figures 3A & S2F) and that RM CAB upregulate MHC-II and costimulatory molecule expression (Figure 3B & 3C). Together, these initial studies showed that *Ct* stimulates nonspecific B cell activation and proliferation in mice, humans, and RM, but compared to murine B cells, human and RM B cells are respectively 3 and 5 orders of magnitude less responsive to this stimulation.

**Figure 2.**
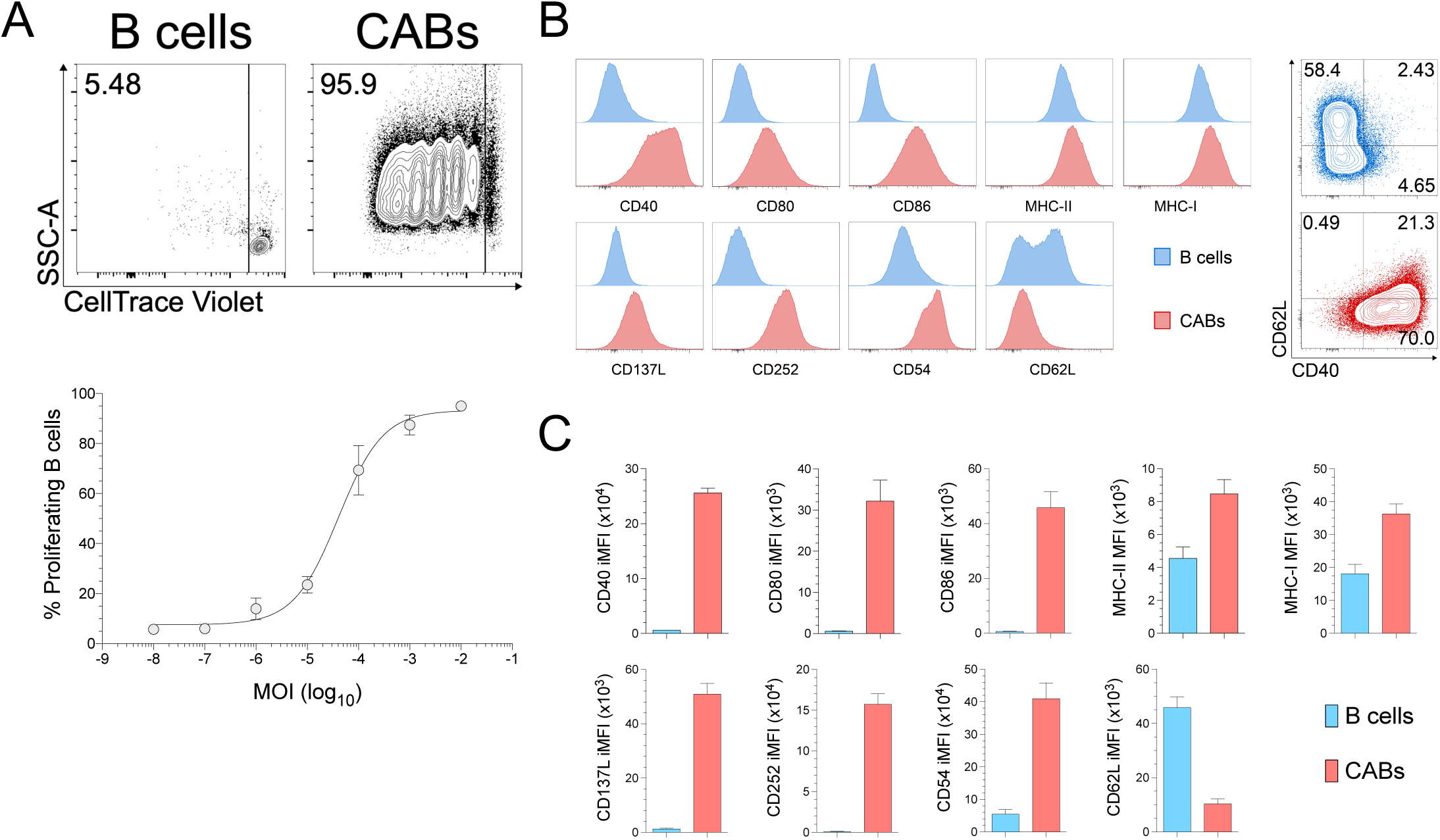
*Ct* nonspecifically stimulates mouse B cells. A) Fluorescently-tagged splenocytes from naïve B6 mice were incubated with inactivated *Ct* L2 elementary bodies at indicated MOI for 4 days and B cell proliferation analyzed by flow cytometry. Top panel displays representative contour plots of fluorescent tracer dilution in B cells (i.e., live CD90^neg^B220^+^ cells) incubated with vehicle (B cells) vs. *Ct* (CAB). Lower panel shows *Ct* dose-responsive B cell proliferation (mean ±SD); maximal response was achieved with lower *Ct* MOI than required by human CAB (Figure 1A). B) Representative histograms and contour plots typify the expression of costimulatory and adhesion molecules by B cells (blue) vs. CAB (red) (quadrant numbers denote percentages). C) Quantified relative expression of these molecules in B cells vs. CABs. Data shown are from 3-7 independent experiments and between-group differences in panel D performed using unpaired Student’s tests; bars denote mean ± SD.

**Figure 3.**
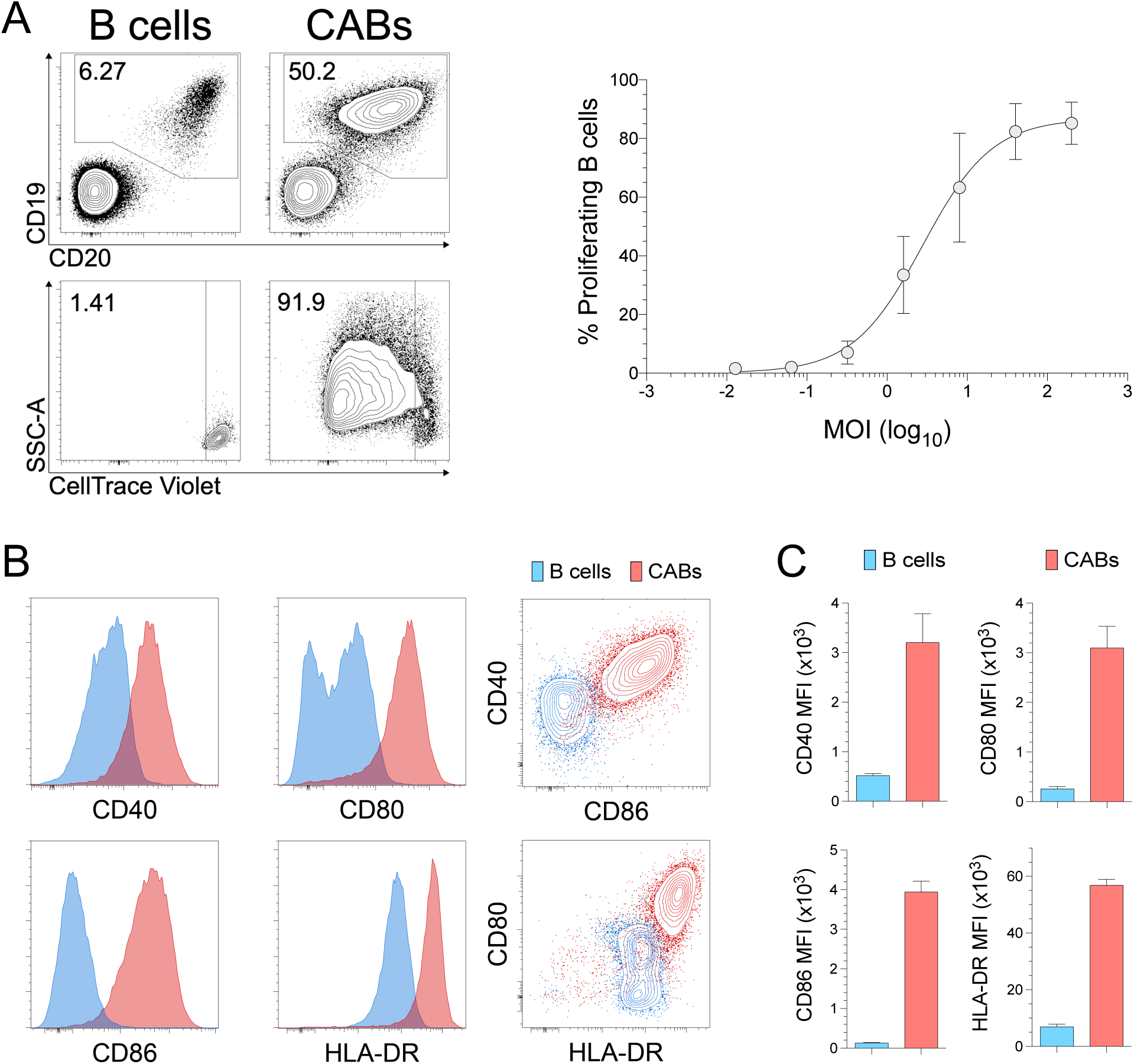
*Ct* nonspecifically activates nonhuman primate B cells. Fluorescently-labeled PBMC from female rhesus macaques (RM) tested seronegative for *Ct*-specific antibodies were incubated for 4 days with varying MOI of inactivated *Ct* L2 EB. A) Quantifying B cell proliferation by flow cytometry, top left panels depict gating strategy used to define B cells (live CD19^+^CD20^+^ cells); bottom left panels depict representative contour plots of fluorescent tracer dilution in B cells incubated with vehicle (B cells) vs. *Ct* (CAB). Right panel displays dose-response curve for B cell proliferation induced by *Ct* (mean ±SD) (MOI needed for maximal response higher in RM vs. mice seen in Figure 1A). B) Representative histograms and contour plots depict expression of costimulatory molecules CD40, CD80, and CD86 and HLA-DR by B cells (blue) vs. CAB (red). C) Quantified relative expression of molecules shown in panel B. Data from 3 separate experiments, between-group differences in panel C were made using unpaired Student’s tests; bars indicate mean ± SD.

Guided by prior report that B cells express multiple Toll like receptors (TLR) and proliferate in response to TLR engagement (17), we used inactivated *Ct* L2 EB in a human TLR (hTLR) ligand screening assay to define Ct engagement of B cell TLR (Figure S2G). This assay identified that *Ct* elicits robust hTLR2 and weak hTLR5 B cell signaling (Figure 4A). Consistent with the lack of TLR5 expression in human B cells (51), we also identified that neutralization of TLR2 (but not TLR5) suppressed *Ct*-induced B cell proliferation (Figure 4B). Other studies demonstrated that inhibition of MyD88 dimerization with the peptidomimetic compound ST2825 also dampened *Ct*-induced human B cell proliferation (Figure 4C) and that *Ct* fails to induce proliferation in splenic B cells from MyD88^-/-^ mice (Figure 4D). *In vitro* stimulation assays also showed that human and mouse B cells do not proliferate when inactivated *Ct* L2 EB is heated to 56°C or proteolytically digested prior to use. These results suggested that chlamydial proteins promote B cell activation (Figure S3A). Because *Chlamydia* MOMP, the immunogenic and immunoaccessible protein that comprises roughly 60% of the total outer membrane protein mass (52), is an adhesin (53) and weak TLR2 agonist (54), we postulated MOMP is a protein that plays a role in *Ct*-induced B cell activation. As predicted, while inactivated *Ct* L2 EB failed to stimulate human B cell proliferation in cultures pre-incubated with polyclonal anti-MOMP antibody or two different anti-MOMP mAb, pre-incubation of EB with anti-*Chlamydia* LPS mAb did not diminish the ability of *Ct* to induce B cell proliferation (Figure 4E). As MOMP and LPS are both present at high density on the surface of *Ct*, these results implied the 3 different anti-MOMP Ab did not inhibit B cell proliferation by sterically hindering bacterium engagement or binding of the B cell Fcγ IIB (FcγRIIB) receptor (55, 56). Our results also showed that while *Ct*-induced proliferation of human B cells depends on TLR2 engagement, unlike mouse B cells (57), TLR2 agonist alone is not sufficient for robust human B cell proliferation (42) (Figure S3B).

**Figure 4.**
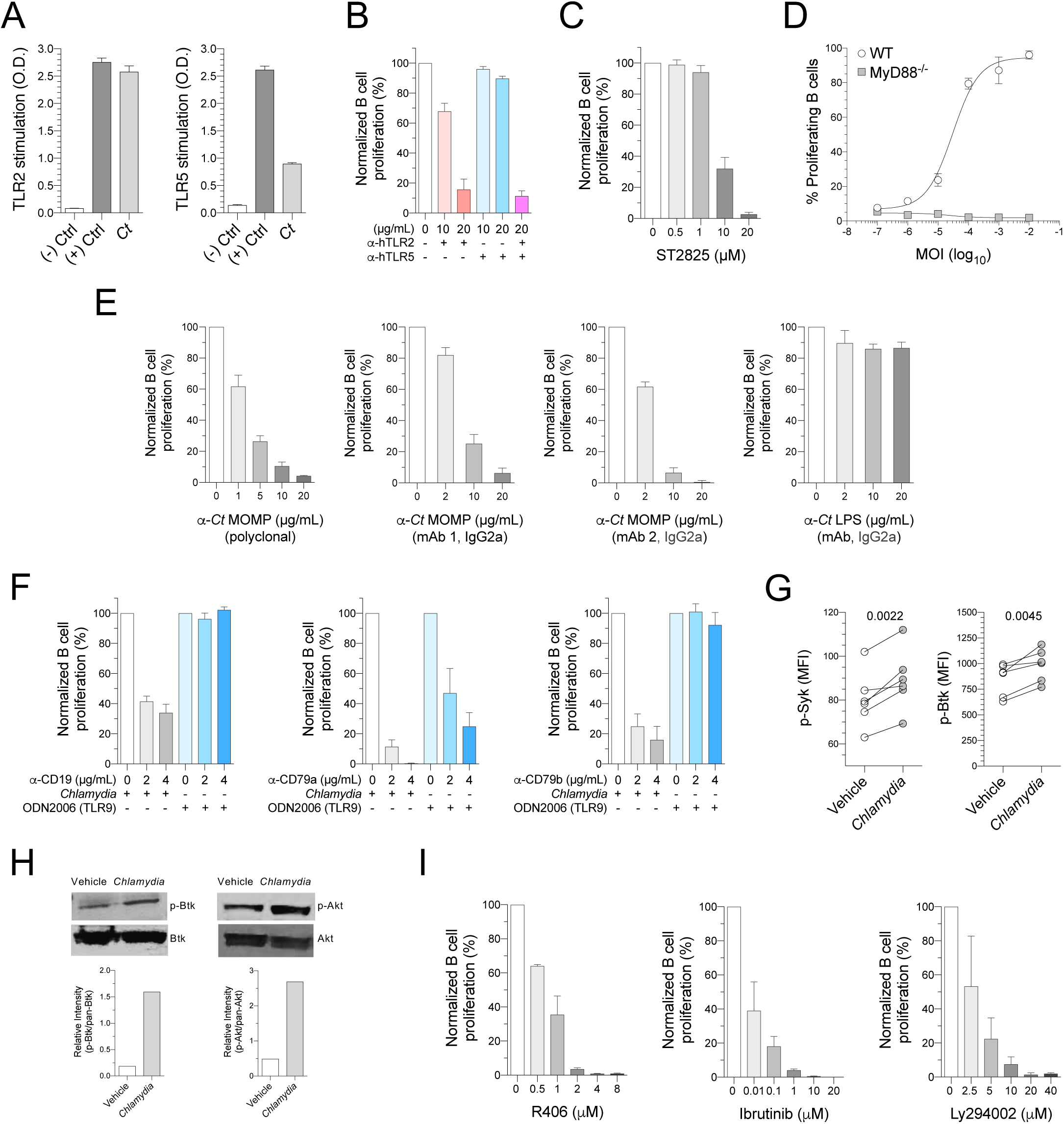
*Ct*-induced human B cell activation requires BCR and TLR signaling. A) Panel shows inactivated *Ct* L2 EB-elicited stimulation of hTLR2 and hTLR5 in HEK293 cells expressing indicated hTLR as assessed by relative NF-κB activation. The MOI of *Ct* L2 EBs used in human B cell studies was the EC50 dose of employed stock. Fluorescently-labeled human PBMC were pre-incubated with B) blocking anti-TLR2 (clone B4H2) and anti-TLR5 (clone Q2G4) mAbs or C) ST2825 (MyD88 dimerization inhibitor) at denoted concentrations for 30 minutes before adding inactivated *Ct* L2 EB. PBMC were cultured for 4 days, and B cell proliferation assessed by flow cytometry. D) Fluorescently-tagged splenocytes from wild type or MyD88^-/-^ mice were incubated for 4 days with indicated MOI of inactivated *Ct* L2 EBs and B cell proliferation evaluated by flow cytometry (mean ± SD). E) Inactivated *Ct* L2 EB were incubated for 30 minutes with polyclonal anti-*Ct* MOMP antibody, 2 anti-*Ct* MOMP mAb (mAb 1, clone HT10 or mAb 2, clone 1297/143), or anti-*Ct* LPS mAb (clone CL21-335.2.3) at specified concentrations before used to stimulate fluorescently-labeled human PBMC. Cultures were incubated for 4 days and proliferation of B cells assessed as in B). F) Fluorescently-labeled human PBMC were pre-incubated for 30 minutes with blocking anti-CD19 (clone HIB19), anti-CD79a (clone JCB117), and anti-CD79b (clone CB3-1) mAb at indicated concentrations before addition of inactivated *Ct* L2 EB or ODN2006 (TLR9 agonist). B cell proliferation was evaluated as in (B). (G) In separate study, human PBMC were stimulated for 15 minutes with *Ct* or vehicle and processed to evaluate phosphorylation of Syk and Btk by phospho flow. H) Human B cells (purified from PBMC by negative immunomagnetic selection) were stimulated with *Ct* or vehicle for 15 minutes and western blot assessed the phosphorylation of Btk and Akt. I) Evaluation of *Ct*-induced B cell proliferation in human PBMC pre-incubated for 30 minutes with R406 (Syk inhibitor), Ibrutinib (Btk inhibitor) or Ly294002 (PI3K inhibitor) at denoted concentrations prior for assessment of B cell activation as described in panel B). Data shown from 3-6 independent experiments; paired Student’s test used to compare paired samples displayed in panel G; bars indicate mean ± SD.

Because TLR signaling and BCR engagement were shown to regulate B cell activation (17, 58), we explored the role of BCR complex proteins in *Ct*-induced human B cell activation. As B cells are non-specifically activated by *Ct* (Figure 1), *Ct* may engage BCR sites not directly related to Ag recognition. We tested this possibility by exploring *Ct*-induced B cell activation in human PBMC pre-incubated with variable concentrations of mAb targeting extracellular portions of CD19 (59), CD79a (60) and CD79b (60, 61). For positive controls, we also incubated PBMC with a TLR9 agonist known to induce B cell proliferation. This study showed that CD19 and CD79b blockade inhibits B cell proliferation induced by *Ct* but not a TLR9 agonist (Figure 4F), whereas anti-CD79a mAb hindered *Ct* and the TLR9 agonist from inducing B cell proliferation. Further supporting a role for the BCR complex in *Ct*-induced B cell activation, we also showed that *Ct* induces kinase phosphorylation critical for BCR signaling, including Syk, Btk and Akt (Figure 4G and 4H) (62–64), while Syk, Btk and PI3K inhibitors, but not ERK (i.e., MAP kinase) inhibition, suppress *Ct*-induced B cell proliferation (Figures 4I and S3C). Viewed together, our initial series of experiments identified that *Ct* nonspecifically activates B cells via TLR signaling and engagement of non-Ag binding sites on the BCR.

### CAB prime antigen-specific T cell immunity

As *Ct* stimulation produced robust and uniform increase in B cell expression of both MHC and costimulatory molecules (Figures 2 & 3), it seemed possible CAB can provide an APC source readily expanded from peripheral blood. In initial work exploring this possibility, we found that mouse and human CAB promote allogeneic naïve CD4^+^ and CD8^+^ T cell proliferation (Figures 5A & 5B) and that human CAB loaded with a pool of viral peptides increase proliferation and effector function of Ag-specific memory CD8^+^ T cells (Figure 5C). Compared to resting B cells, mouse CAB loaded with fluorescently-conjugated OVA or a self-quenched OVA conjugate (DQ-OVA) that fluoresces upon proteolytic degradation more readily processed exogenous soluble antigen (Figures 5D & 5E). As CAB increased expression of adhesion molecules that promote APC trafficking to secondary lymphoid organs (Figure 2), we also explored the ability of CAB to prime CD8^+^ T cell immunity *in vivo*. For this study, fluorescently CTV-labeled OVA-specific TCR transgenic CD8^+^ T cells were transferred from OT-I mice to wild type B6 mice. The next day, OVA-loaded or unloaded syngeneic CAB were transferred to these recipients. Of note, no further purification steps were needed prior to transfer as final CAB suspensions contained >95% CABs and no DC (Figure S2B & S2E). Congruent with our *in vitro* data, OVA-loaded CAB elicited robust proliferation of Ag-specific CD8^+^ T cells (Figure 5F). Other *in vivo* studies showed CAB loaded with the immunodominant OVA epitope (OVA257-264) were more potent activators of OT-I cells than OVA-loaded CAB, while OVA-loaded resting B cells inefficiently primed CD8^+^ T cell responses (Figure 5G). Performing comparable studies with gp100-specific TCR transgenic CD8^+^ T cells and CAB loaded with the immunodominant gp100 peptide (gp10025-33) showed that CAB also induce *in vivo* CD8^+^ T cell responses to tumor-associated Ag (Figure 5H). Comparing freshly OVA-loaded CAB vs. thawed CAB loaded with OVA prior to cryopreservation, we found comparable purity and activation levels (Figure S3D-F) and established that cryopreservation did not diminish the capacity of CAB to induce Ag-specific CD8^+^ T cell proliferation (Figure 5I).

**Figure 5.**
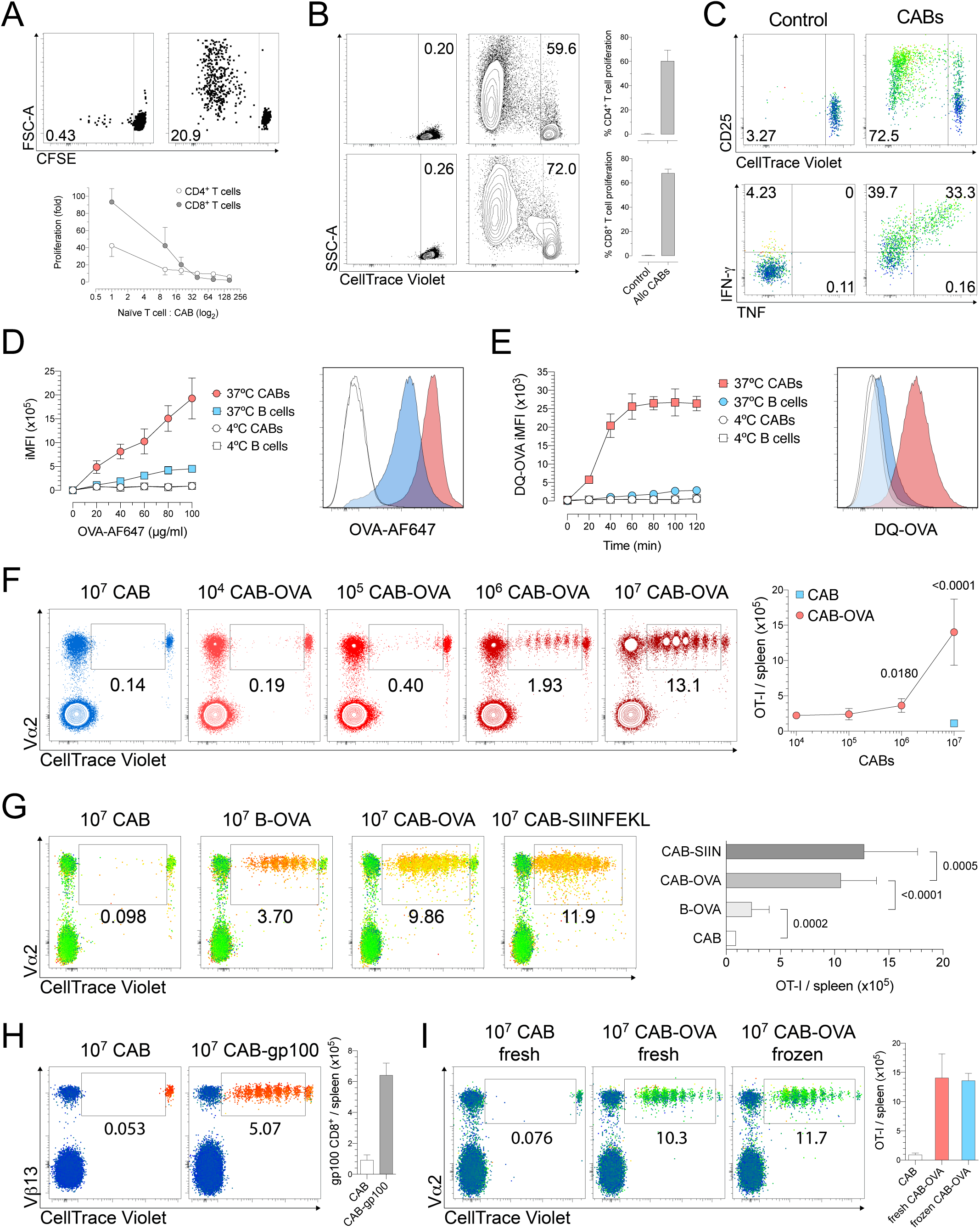
CAB prime CD8^+^ T cell proliferation. A) CAB generated from B6 mice splenocytes were incubated with fluorescently-labeled allogeneic T cells from uninfected Balb/cJ mice for 4 days and CD4^+^ and CD8^+^ T cell proliferation determined by flow cytometry. Top panel depicts representative dot plots, lower panel displays fold increases in proliferating T cells (mean ± SD). B) Human CAB were co-incubated with fluorescently-tagged allogeneic naïve T cells for 7 days and CD4^+^ and CD8^+^ T cell proliferation defined by flow cytometry. Top and lower panels display representative contour plots and quantified proliferation of CD4^+^ and CD8^+^ T cells, respectively (bars indicate mean ± SD). C) Human CAB loaded with a CEF peptide pool were incubated with fluorescently-labeled autologous memory CD8^+^ T cells for 7 days and proliferation (top panels) and cytokine production (lower panels) assessed by flow cytometry (representative heatmap statistical plots shown, color scale indicates SSC levels). Murine CAB or resting B cells were incubated (D) for 1 h with OVA-AF647 (0-120 μg/ml) at 4°C or 37°C or (E) for 0-120 minutes with DQ-OVA (10 μg/ml) at 4°C or 37°C, and fluorescence levels defined by flow cytometry (mean ± SD). F) 10^6^ fluorescently labeled OT-I cells were i.v. transferred to B6 mice. The next day, the indicated numbers of unloaded (CAB) or OVA-loaded CAB were i.v. transferred. Mice were euthanized 3 days later, and spleens processed into single-cell suspensions to define OT-I proliferation by flow cytometry. Left panels display representative contour plots and right panel depicts OT-1 proliferation elicited by unloaded CAB or indicated numbers of OVA-loaded CAB (mean ± SD), comparison between groups that received unloaded CAB vs. OVA-loaded CAB was made using one-way ANOVA and Dunnett’s multiple comparison post hoc test. G) B6 mice transferred OT-I cells as in (F) received OVA-loaded CABs (CAB-OVA), SIINFEKL (OVA257-264)- loaded CABs (CAB-SIINFEKL), OVA-loaded resting B cells (B-OVA), or unloaded CAB (CAB). Left panels show representative heatmap statistical plots for OT-1 cell proliferation, color scale denotes CD44 expression levels. Right panel compares OT-I cell proliferation; p<0.0001 for groups that received antigen-loaded CABs vs. unloaded CAB. H) 10^6^ fluorescently-labeled gp-100-specific CD8^+^ T cells were transferred into B6 mice. 24 hours later, 10^7^ CABs loaded with matching immunogenic peptide (gp10025-33) were i.v. administered. Antigen-specific CD8^+^ T cell proliferation was defined 3 days later. Left panel shows representative heatmap statistical plots, color scale denotes CD90.1 expression level. I) In other *in vivo* priming studies performed as in (G), mice were administered unloaded CAB (CAB), freshly prepared CAB loaded with OVA (fresh CAB-OVA), or thawed CAB loaded with OVA prior to cryopreservation (frozen CAB-OVA). Left panel shows representative heatmap statistical plots, color scale denotes forward scatter levels. Right panel shows OT-I cell proliferation, using the unpaired Student’s test to compare proliferation in mice that received fresh CAB-OVA vs. frozen CAB-OVA. Data displayed are from 3-5 independent experiments; in panels G), H) and I), bars indicate mean ± SD.

To extend our findings, PBMC from RM were used to develop methodology for CAB production (Figure S1E) and define the ability of RM CAB to elicit Ag-specific CD8^+^ T cell immunity. For the latter, we loaded CAB overnight with the model Ag keyhole limpet hemocyanin (KLH) and used negative immunomagnetic selection to remove T cells prior to cryopreservation (Figure S4A-H). These autologous KLH-loaded CAB were thawed immediately before administration (over a 6-week period, RM received a total of 3 doses of 5 x 10^7^ KLH-loaded CAB) (Figure 6A). To evaluate Ag-specific T cell responses, PBMC obtained at time points shown in Figure 6A and splenocytes collected at euthanasia (7 weeks after initial CAB injection) were used for ICS. As indicated by the levels of IFN-γ secreted in response to *ex vivo* stimulation with KLH antigen, this study identified that autologous RM CAB effectively promote formation of Ag-specific CD8^+^ T cell responses (primary endpoint) (Figure 6B and C).

**Figure 6.**
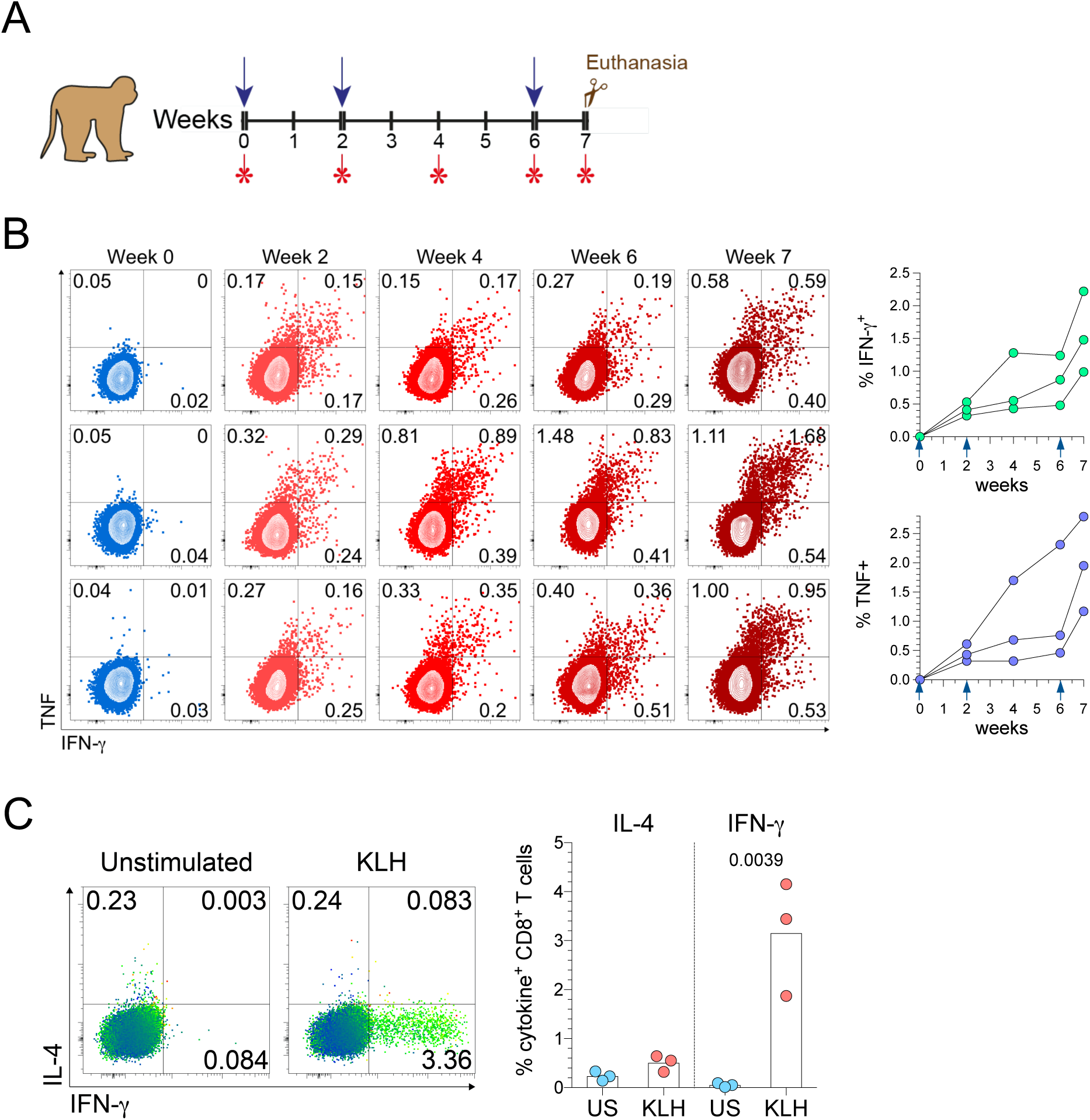
Antigen-loaded CAB prime *in vivo* antigen-specific CD8^+^ T cells responses in nonhuman primates. A) Depiction of study design used to investigate CD8+ T cell immunity elicited in RM (n = 3), blue arrows and red asterisks indicate KLH-loaded CAB injections and blood draws, respectively. B) Cryopreserved PBMC were thawed and incubated overnight with the model antigen keyhole limpet hemocyanin (KLH) (the last 6 hours in presence of BFA). ICS assessed IFN-γ and TNF production by flow cytometry. Left panels show contour plots of CD8^+^ T cell cytokine production by individual RM at indicated time point. Right panels display individual IFN-γ and TNF expression levels. C) Cryopreserved splenocytes were thawed and incubated overnight with KLH, as in (B). Left panels illustrate representative heatmap statistical plots (color scale indicates FSC levels) of IFN-γ and IL-4 production by CD8^+^ T cells, right panels display levels in individual RM (bars indicate means).

### CAB boost *in vivo* Ag-specific T cell immunity

As *ex vivo* studies showed that Ag-loaded CAB generate Ag-specific CD8^+^ T immunity (Figure 6), we also explored the ability of CAB to boost host responses to tumor and microbial Ag. Our initial study used an ocular HSV-2 infection model in which wild type mice received 3 doses of unloaded CAB or CAB loaded with the immunodominant HSV-2 peptide (i.e., gB498-505) prior to corneal infection with a typically lethal inoculum of virus. Whilst all mice injected with unloaded CAB developed fatal encephalopathy, no morbidity or mortality occurred in mice that received virus peptide-loaded CAB prior to infection (Figure 7A). In separate study, murine models of E.G7-OVA and B16-F10 tumor explored the ability of Ag-loaded CAB to prophylactically deter tumor development. In these studies, mice administered CAB loaded with immunodominant peptide or cognate Ag prior to subcutaneous (E.G7-OVA) or intravenous (B16-F10) tumor cell injection displayed attenuated tumor growth or no development of tumor (Figures 7B and 7C). Of note, mice protected from OVA tumor development remained tumor-free when re-challenged with the same tumor cells. In separate study, we used the E.G7-OVA tumor model to assess the effects of Ag-loaded CAB administered after tumor injection. Starting 5 days after wild type mice were subcutaneously injected with 7.5 x 10^5^ E.G7-OVA tumor cells (a time point when tumors were palpable), mice received unloaded CAB or OVA-loaded CAB (treatment was every other day for a total of five doses). Consistent with results from our tumor prophylaxis studies, mice administered OVA-loaded CAB displayed significantly lower tumor burden (primary endpoint) and 40% of treated animals completely rejected tumor (Figures 7D and 7E). OVA-loaded CAB injections were also associated with significantly lower frequencies of splenic MDSC and FoxP3^+^ regulatory T cells (Figures S4I-J, 7F & 7G) (secondary endpoints), whereas CAB-mediated prophylactic and therapeutic effects were eliminated in mice concomitantly administered CD8-depleting mAb. Together, this last set of studies identified that host immune responses generated by CAB vaccination improved study outcomes in mice by enhancing antiviral and antitumor immunity.

**Figure 7.**
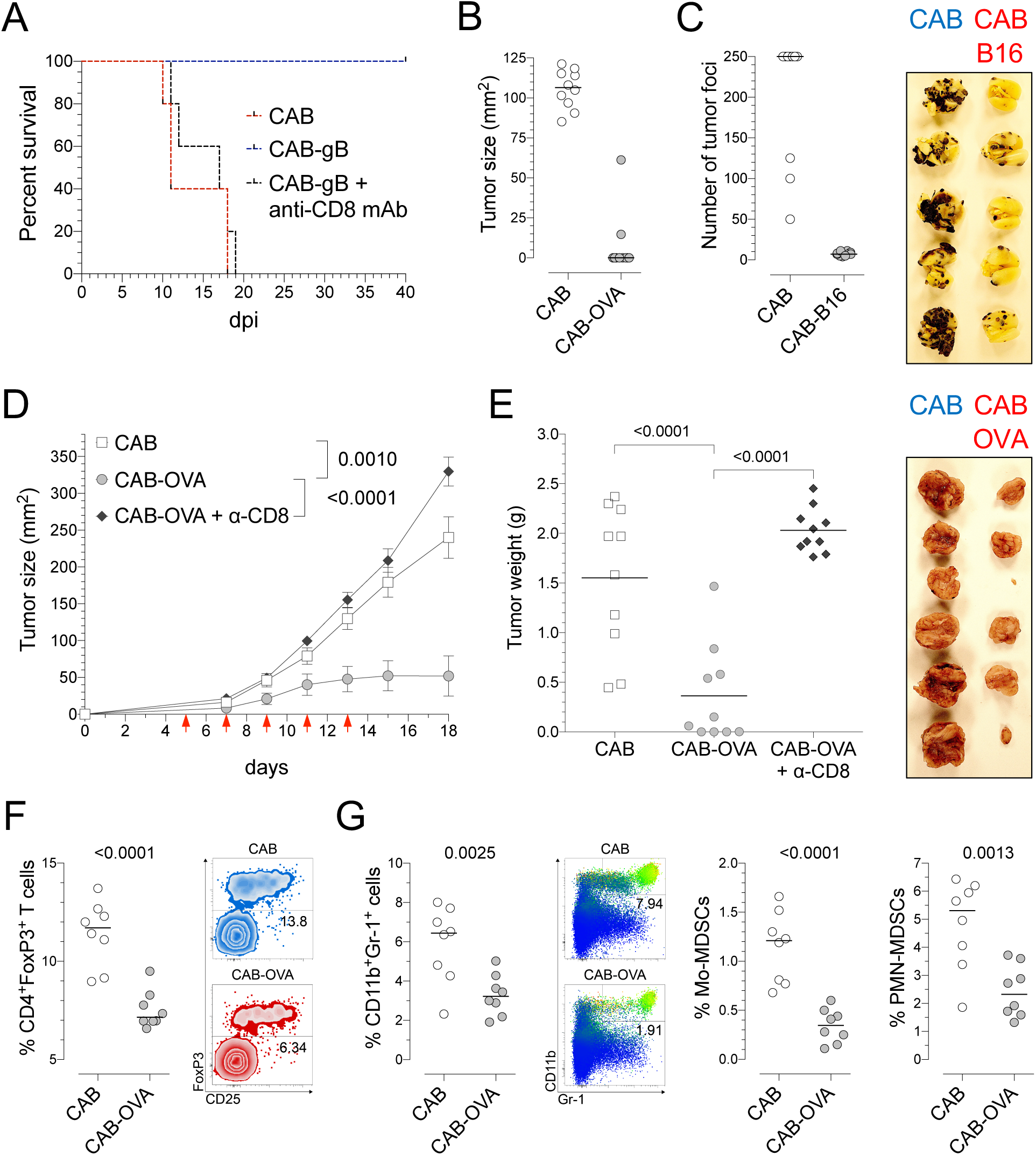
Antigen-loaded murine CAB stimulate *in vivo* antigen-specific CD8^+^ T cell immunity against tumor and virus. A) B6 mice were immunized with 3 doses of unloaded CAB (CAB) or CAB loaded with gB498-505 (CAB-gB). A portion of the latter group also received anti-CD8 depleting mAb (CAB-gB + anti-CD8 mAb) prior to corneal HSV-2 infection (n=5 per group). Mice were monitored daily after infection. Kaplan-Meier survival curve depicts HSV-induced mortality. B) B6 mice were immunized similar to (A) but instead administered unloaded CAB (CAB) or OVA-loaded CAB (CAB-OVA). Mice were s.c. injected with 10^6^ E.G7-OVA tumor cells and tumor size gaged 10 days later (right panel displays representative images). C) B6 mice were immunized as in (A) but administered CAB mock-loaded (CAB) or loaded with TRP-2180-188 and gp10025-33 (CAB-B16). 18 days after i.v. administration of 1.5 x 10^5^ B16-F10 tumor cells, mice were euthanized to enumerate metastatic lung foci (right panel shows representative images). D) B6 mice were s.c. injected with 7.5 x 10^5^ E.G7-OVA tumor cells and 5 days later (when tumors were palpable) were i.v. administered 10^7^ unloaded CAB (CAB) or OVA-loaded CAB (CAB-OVA) every other day (total of 5 doses). A third group of tumor-injected mice (n=10 per group) received an identical course of OVA-loaded CAB and anti-CD8 depleting mAb (CAB- OVA + α-CD8). Tumor growth (mean ± SEM) was calculated every other day to allow between-group comparison of growth. E) In other studies with mice treated as in (D), tumor weights were measured 18 days after tumor injection; right panel shows representative images. Frequencies of splenic F) FoxP3^+^ regulatory T cells and G) MDSC (i.e., CD11b^+^Gr-1^+^ cells, Mo-MDSC and PMN-MDSC) in mice administered unloaded CAB (CAB) vs. OVA-loaded CAB (CAB-OVA) were defined by flow cytometry. G) Representative heatmap statistical plots depict CD11b and Gr-1 expression in total splenocytes, gate defines total CD11b^+^Gr-1^+^ MDSCs, color scale indicates levels of side scatter. Data are from 2-3 independent experiments. Between-group differences in panels B, C, E, F, and G were made using unpaired Student’s tests.

## DISCUSSION

While not the primary objective of this investigation, current results offer insight into the human host response to chlamydia. Genital *Ct* infection in women is usually asymptomatic and slowly eradicated (29), a clinical presentation that suggests *Ct* evades host defenses to establish chronic subclinical infection. *Ct*-infected human endometrial tissue is also characterized by B cell-rich cellular aggregates and raised expression of genes associated with B cell development, including CD40, CD79a, CD79b, CD86, RAG1, and the heavy and light immunoglobulin chains (24–26). Combining past observation with current results, we speculate *Ct* evolved mechanisms to elicit non-specific B cell activation and delay formation of *Ct*-specific immune responses. This possibility is consistent with other pathogens shown to promote nonspecific antibody production to subvert development of effective host immunity (20, 65). Current identification of *Ct*-induced non-specific B cell activation therefore identifies a potential mechanism for immunoevasion that informs pathogenesis of genital *Ct* infection and may contribute to *Ct* vaccine design.

However, the main goals for this investigation were to explore both mechanisms of *Chlamydia*-mediated B cell activation and the ability CAB to function as APC. Addressing the first objective, we found that nonspecific human B cell activation is induced by interactions between chlamydial MOMP and B cell TLR2, CD19, and BCR CD79a/b complexes. We also identified that CD79a blockade dampens human B cell proliferation induced by *Ct* or a TLR9 agonist and that CD19 and CD79b blockade diminishes *Ct*-induced but not TLR9-induced B cell proliferation. While novel, these results are consistent with prior work in mice in which CD79a blockade dampened B cell proliferation induced by TLR and BCR signals and BCR crosslinking stimulated weak B cell proliferation (66, 67). We also show that BCR crosslinking alone does not stimulate human B cell proliferation (Figure S3B), results that imply this proliferation is potentiated by MOMP-mediated BCR signals that sensitize cells to TLR2 ligand binding (42). This possibility is congruent with mechanisms that *Moraxella catarrhalis* and *S. aureus* use to stimulate T cell-independent B cell proliferation (68, 69). While MOMP is a known weak TLR2 agonist (54, 70), future studies will need to explore the possibility that other chlamydial proteins or LPS trigger TLR2 signaling in human B cells (71, 72). Further study is also needed to define the differential sensitivity to *Ct*-induced activation in B cells from mice, NHP, and humans (Figures 2 & 3), an observation feasibly related to dissimilar TLR expression in B cells from these species (73, 74).

Addressing the second objective (i.e., the ability of CAB to function as APC), our initial studies showed that CAB uniformly upregulate costimulatory molecules that induce and maintain T cell immunity, including CD40, CD80, CD86, CD137L, and CD252 (75–77). Compared to resting B cells, human CAB also significantly increase expression of CD54, CD58, and CD62L, adhesion molecules that promote T cell immunity *in vivo* (78). In contrast to higher CD62L levels in CAB vs. resting B cells from human PBMC, expression of this adhesion molecule is higher in resting B cells vs. CAB generated from murine splenocytes. Despite these lower levels of CD62L, our follow-up studies identified that murine CAB effectively prime *in vivo* CD8^+^ T cell immunity. As lower CD62L levels were shown to impair trafficking of DC to secondary lymphoid organs (79), our results imply a lesser role for CD62L in CAB trafficking or greater contribution from other adhesion molecules or chemokine receptors (e.g., CCR7 (78). In addition, the current study also shows *in vitro* B cell stimulation with *Ct* does not promote plasma cell formation. This response is contrary the differentiation of plasma cells induced by *in vivo* infection of humans with *Ct* and mouse with *Cm* (23, 24, 80), and potentially related to the *in vitro* absence of IL-21, CD154, and other microenvironment signals that stimulate plasma cell generation *in vivo* (81).

While defining the signals required for *Ct*-induced plasma cell differentiation and mechanisms of CAB trafficking *in vivo* awaits further research, current results already suggest that CAB may provide basis for a vaccine platform that elicits robust Ag-specific CD8^+^ T cell immunity against tumors and microbial pathogens. B cells are the largest APC pool in peripheral blood and ready expansion of large numbers of CAB may offer advantages over DC-based immunotherapy (3). While *in vitro* study indicated that Ag-loaded DC prime cytotoxic T lymphocyte (CTL) immunity (82), no significant improvement in survival was achieved in clinical trials that administered DC-based vaccines to individuals with advanced prostate cancer (83) or melanoma (84). DC-based vaccines likewise display ineffective trafficking, and even after intranodal administration, < 5% of mature DC reached lymph node T cell zones (85). While intralymphatic delivery may enhance effective DC trafficking, clinical complications are associated with intralymphatic port placement (86). It is also difficult to generate adequate numbers of uniformly activated DC that maintain APC function after cryopreservation (6–11, 87). Compared to limitations currently associated with DC-based vaccines, our investigation identifies that *Chlamydia*-induced *ex vivo* stimulation of B cells produces large numbers of uniformly activated APC, and this ability to readily expand the most plentiful pool of APC in peripheral blood may facilitate more frequent administrations with larger APC numbers. Of note, these parameters were associated with enhanced antitumor immunity and improved clinical outcomes in studies that employed DC-based platforms (3, 6, 7).

Congruent with the ability of CAB to function as APC, our study identifies that CAB effectively enhance Ag-specific CD8^+^ T cell immunity *in vivo*. Specifically, intravenous injection of RM with KLH-loaded CAB RM enhanced Ag-specific CD8^+^ T cell effector function and mice injected with cognate Ag-loaded CAB generated CD8^+^ T cell immune responses that protected against virus infection and produced robust prophylactic and therapeutic antitumor activity. Although direct comparisons were not performed in the current investigation, our results imply that more robust antitumor activity is elicited by administration of CAB vs. CD40-activated B cells (CD40B) (88).

However, these prior explorations intraperitoneally injected mice with CD40B, an administration route that decreases CD40B trafficking to secondary lymphoid organs and may have lessened the ability of this B cell-based vaccine platform to boost antitumor activity (89, 90). In addition to the robust *in vivo* antitumor and antiviral immunity generated by CAB, current results indicate that cryopreservation does not diminish the ability of CAB to induce *in vivo* CTL immunity. For APC-based vaccines, the ability to use cryopreserved cells seems likely to simplify treatment logistics and expand delivery options. As another potential benefit of a CAB-based vaccine, compared to controls, spleens from mice administered Ag-loaded CAB contained significantly fewer FoxP3^+^ regulatory T cells and MDSC, immune cells shown to impair efficacy of cancer immunotherapy (91, 92). Taken together, current results suggest it will be possible to leverage *Ct*-induced B cell activation into development of a vaccine platform that significantly boosts Ag-specific T cell immunity. Ongoing research in our lab towards this goal include defining CAB trafficking patterns, molecular mechanisms by which CAB prime T cell immunity, and *in vivo* durability of CAB-induced T cell memory.

## ACKNOWLEDGMENTS

Authors acknowledge technical expertise provided by Ao Xie, Kevin Henschel, and Mohammed Ghonime and the support provided by the OSU Comprehensive Cancer Center and the OSU Analytical Cytometry Shared Resource (supported by NCI Grant P30 CA016058).

## FOOTNOTES

### Author contributions

RDVM and TLC conceived this research, supervised studies, analyzed data, and composed the initial manuscript draft. RDVM, JGC, SF and NEQC performed experiments. DJC directed and performed the RM-related research. All authors were involved with manuscript preparation.

### Data availability

All data are available in the main text or the supplementary materials upon reasonable request.

## REFERENCES

1. Santos PM, Butterfield LH. Dendritic Cell-Based Cancer Vaccines. Journal of immunology (Baltimore, Md : 1950). 2018;200(2):443–9.

2. Le DT, Pardoll DM, Jaffee EM. Cellular vaccine approaches. Cancer journal (Sudbury, Mass). 2010;16(4):304–10.

3. Anguille S, Smits EL, Lion E, van Tendeloo VF, Berneman ZN. Clinical use of dendritic cells for cancer therapy. The Lancet Oncology. 2014;15(7):e257–67.

4. Harari A, Graciotti M, Bassani-Sternberg M, Kandalaft LE. Antitumour dendritic cell vaccination in a priming and boosting approach. Nat Rev Drug Discov. 2020;19(9):635–52.

5. Wculek SK, Cueto FJ, Mujal AM, Melero I, Krummel MF, Sancho D. Dendritic cells in cancer immunology and immunotherapy. Nature reviews Immunology. 2020;20(1):7–24.

6. Draube A, Klein-Gonzalez N, Mattheus S, Brillant C, Hellmich M, Engert A, et al. Dendritic cell based tumor vaccination in prostate and renal cell cancer: a systematic review and meta-analysis. PloS one. 2011;6(4):e18801.

7. MartIn-Fontecha A, Sebastiani S, Hopken UE, Uguccioni M, Lipp M, Lanzavecchia A, et al. Regulation of dendritic cell migration to the draining lymph node: impact on T lymphocyte traffic and priming. The Journal of experimental medicine. 2003;198(4):615–21.

8. Hayden H, Friedl J, Dettke M, Sachet M, Hassler M, Dubsky P, et al. Cryopreservation of monocytes is superior to cryopreservation of immature or semi-mature dendritic cells for dendritic cell-based immunotherapy. Journal of immunotherapy (Hagerstown, Md : 1997). 2009;32(6):638–54.

9. Buhl T, Legler TJ, Rosenberger A, Schardt A, Schon MP, Haenssle HA. Controlled-rate freezer cryopreservation of highly concentrated peripheral blood mononuclear cells results in higher cell yields and superior autologous T-cell stimulation for dendritic cell-based immunotherapy. Cancer immunology, immunotherapy : CII. 2012;61(11):2021–31.

10. Zhou Q, Zhang Y, Zhao M, Wang X, Ma C, Jiang X, et al. Mature dendritic cell derived from cryopreserved immature dendritic cell shows impaired homing ability and reduced anti-viral therapeutic effects. Scientific reports. 2016;6:39071.

11. Chung DJ, Romano E, Pronschinske KB, Shyer JA, Mennecozzi M, St Angelo ET, et al. Langerhans-type and monocyte-derived human dendritic cells have different susceptibilities to mRNA electroporation with distinct effects on maturation and activation: implications for immunogenicity in dendritic cell-based immunotherapy. Journal of translational medicine. 2013;11:166.

12. Wennhold K, Shimabukuro-Vornhagen A, Theurich S, von Bergwelt-Baildon M. CD40-activated B cells as antigen-presenting cells: the final sprint toward clinical application. Expert review of vaccines. 2013;12(6):631–7.

13. Tretter T, Venigalla RK, Eckstein V, Saffrich R, Sertel S, Ho AD, et al. Induction of CD4+ T-cell anergy and apoptosis by activated human B cells. Blood. 2008;112(12):4555–64.

14. Tu W, Lau YL, Zheng J, Liu Y, Chan PL, Mao H, et al. Efficient generation of human alloantigen-specific CD4+ regulatory T cells from naive precursors by CD40-activated B cells. Blood. 2008;112(6):2554–62.

15. Kleindienst P, Brocker T. Concerted antigen presentation by dendritic cells and B cells is necessary for optimal CD4 T-cell immunity in vivo. Immunology. 2005;115(4):556–64.

16. Parekh VV, Prasad DV, Banerjee PP, Joshi BN, Kumar A, Mishra GC. B cells activated by lipopolysaccharide, but not by anti-Ig and anti-CD40 antibody, induce anergy in CD8+ T cells: role of TGF-beta 1. Journal of immunology (Baltimore, Md : 1950). 2003;170(12):5897–911.

17. Hua Z, Hou B. TLR signaling in B-cell development and activation. Cellular & molecular immunology. 2013;10(2):103–6.

18. Lapointe R, Bellemare-Pelletier A, Housseau F, Thibodeau J, Hwu P. CD40-stimulated B lymphocytes pulsed with tumor antigens are effective antigen-presenting cells that can generate specific T cells. Cancer research. 2003;63(11):2836–43.

19. Shimabukuro-Vornhagen A, Kondo E, Liebig T, von Bergwelt-Baildon M. Activated human B cells: stimulatory or tolerogenic antigen-presenting cells? Blood. 114. United States2009. p. 746–7; author reply 7.

20. Nothelfer K, Sansonetti PJ, Phalipon A. Pathogen manipulation of B cells: the best defence is a good offence. Nature reviews Microbiology. 2015;13(3):173–84.

21. Levitt D, Barol J. The immunobiology of Chlamydia. Immunology today. 1987;8(7-8):246–51.

22. Levitt D, Newcomb RW, Beem MO. Excessive numbers and activity of peripheral blood B cells in infants with Chlamydia trachomatis pneumonia. Clinical immunology and immunopathology. 1983;29(3):424–32.

23. Vicetti Miguel RD, Quispe Calla NE, Cherpes TL. Setting Sights on Chlamydia Immunity’s Central Paradigm: Can We Hit a Moving Target? Infection and immunity. 2017;85(7).

24. Vicetti Miguel RD, Harvey SA, LaFramboise WA, Reighard SD, Matthews DB, Cherpes TL. Human female genital tract infection by the obligate intracellular bacterium Chlamydia trachomatis elicits robust Type 2 immunity. PloS one. 2013;8(3):e58565.

25. Vicetti Miguel RD, Reighard SD, Chavez JM, Rabe LK, Maryak SA, Wiesenfeld HC, et al. Transient detection of Chlamydial-specific Th1 memory cells in the peripheral circulation of women with history of Chlamydia trachomatis genital tract infection. American journal of reproductive immunology (New York, NY : 1989). 2012;68(6):499–506.

26. Reighard SD, Sweet RL, Vicetti Miguel C, Vicetti Miguel RD, Chivukula M, Krishnamurti U, et al. Endometrial leukocyte subpopulations associated with Chlamydia trachomatis, Neisseria gonorrhoeae, and Trichomonas vaginalis genital tract infection. American journal of obstetrics and gynecology. 2011;205(4):324.e1-7.

27. Chavez JM, Vicetti Miguel RD, Cherpes TL. Chlamydia trachomatis infection control programs: lessons learned and implications for vaccine development. Infectious diseases in obstetrics and gynecology. 2011;2011:754060.

28. Vicetti Miguel RD, Cherpes TL. Hypothesis: Chlamydia trachomatis infection of the female genital tract is controlled by Type 2 immunity. Medical hypotheses. 2012;79(6):713–6.

29. Batteiger BE, Xu F, Johnson RE, Rekart ML. Protective immunity to Chlamydia trachomatis genital infection: evidence from human studies. The Journal of infectious diseases. 2010;201 Suppl 2:S178–89.

30. Hou B, Reizis B, DeFranco AL. Toll-like receptors activate innate and adaptive immunity by using dendritic cell-intrinsic and -extrinsic mechanisms. Immunity. 2008;29(2):272–82.

31. Overwijk WW, Restifo NP. B16 as a mouse model for human melanoma. Current protocols in immunology. 2001;Chapter 20:Unit 20.1.

32. Vicetti Miguel RD, Cherpes TL, Watson LJ, McKenna KC. CTL induction of tumoricidal nitric oxide production by intratumoral macrophages is critical for tumor elimination. Journal of immunology (Baltimore, Md : 1950). 2010;185(11):6706–18.

33. Cherpes TL, Harvey SA, Phillips JM, Vicetti Miguel RD, Melan MA, Quispe Calla NE, et al. Use of transcriptional profiling to delineate the initial response of mice to intravaginal herpes simplex virus type 2 infection. Viral immunology. 2013;26(3):172–9.

34. Vicetti Miguel RD, Quispe Calla NE, Dixon D, Foster RA, Gambotto A, Pavelko SD, et al. IL-4-secreting eosinophils promote endometrial stromal cell proliferation and prevent Chlamydia-induced upper genital tract damage. Proceedings of the National Academy of Sciences of the United States of America. 2017;114(33):E6892–e901.

35. Vicetti Miguel RD, Quispe Calla NE, Pavelko SD, Cherpes TL. Intravaginal Chlamydia trachomatis Challenge Infection Elicits TH1 and TH17 Immune Responses in Mice That Promote Pathogen Clearance and Genital Tract Damage. PloS one. 2016;11(9):e0162445.

36. Vicetti Miguel RD, Henschel KJ, Duenas Lopez FC, Quispe Calla NE, Cherpes TL. Fluorescent labeling reliably identifies Chlamydia trachomatis in living human endometrial cells and rapidly and accurately quantifies chlamydial inclusion forming units. Journal of microbiological methods. 2015;119:79–82.

37. Vicetti Miguel RD, Maryak SA, Cherpes TL. Brefeldin A, but not monensin, enables flow cytometric detection of interleukin-4 within peripheral T cells responding to ex vivo stimulation with Chlamydia trachomatis. Journal of immunological methods. 2012;384(1-2):191–5.

38. Vicetti Miguel RD, Hendricks RL, Aguirre AJ, Melan MA, Harvey SA, Terry-Allison T, et al. Dendritic cell activation and memory cell development are impaired among mice administered medroxyprogesterone acetate prior to mucosal herpes simplex virus type 1 infection. Journal of immunology (Baltimore, Md : 1950). 2012;189(7):3449–61.

39. Quispe Calla NE, Ghonime MG, Cherpes TL, Vicetti Miguel RD. Medroxyprogesterone acetate impairs human dendritic cell activation and function. Human reproduction (Oxford, England). 2015;30(5):1169–77.

40. Quispe Calla NE, Vicetti Miguel RD, Mei A, Fan S, Gilmore JR, Cherpes TL. Dendritic cell function and pathogen-specific T cell immunity are inhibited in mice administered levonorgestrel prior to intranasal Chlamydia trachomatis infection. Scientific reports. 2016;6:37723.

41. Bielinska AU, Makidon PE, Janczak KW, Blanco LP, Swanson B, Smith DM, et al. Distinct pathways of humoral and cellular immunity induced with the mucosal administration of a nanoemulsion adjuvant. Journal of immunology (Baltimore, Md : 1950). 2014;192(6):2722–33.

42. Bekeredjian-Ding I, Foermer S, Kirschning CJ, Parcina M, Heeg K. Poke weed mitogen requires Toll-like receptor ligands for proliferative activity in human and murine B lymphocytes. PLoS One. 2012;7(1):e29806.

43. Liu TM, Ling Y, Woyach JA, Beckwith K, Yeh YY, Hertlein E, et al. OSU-T315: a novel targeted therapeutic that antagonizes AKT membrane localization and activation of chronic lymphocytic leukemia cells. Blood. 2015;125(2):284–95.

44. Irish JM, Czerwinski DK, Nolan GP, Levy R. Kinetics of B cell receptor signaling in human B cell subsets mapped by phosphospecific flow cytometry. Journal of immunology (Baltimore, Md : 1950). 2006;177(3):1581–9.

45. Bronte V, Brandau S, Chen SH, Colombo MP, Frey AB, Greten TF, et al. Recommendations for myeloid-derived suppressor cell nomenclature and characterization standards. Nature communications. 2016;7:12150.

46. Faul F, Erdfelder E, Lang AG, Buchner A. G*Power 3: a flexible statistical power analysis program for the social, behavioral, and biomedical sciences. Behav Res Methods. 2007;39(2):175–91.

47. Faul F, Erdfelder E, Buchner A, Lang AG. Statistical power analyses using G*Power 3.1: tests for correlation and regression analyses. Behav Res Methods. 2009;41(4):1149–60.

48. Bard J, Levitt D. Chlamydia trachomatis stimulates human peripheral blood B lymphocytes to proliferate and secrete polyclonal immunoglobulins in vitro. Infection and immunity. 1984;43(1):84–92.

49. Levitt D, Danen R, Bard J. Both species of chlamydia and two biovars of Chlamydia trachomatis stimulate mouse B lymphocytes. Journal of immunology (Baltimore, Md : 1950). 1986;136(11):4249–54.

50. Bard J, Levitt D. Chlamydia trachomatis (L2 serovar) binds to distinct subpopulations of human peripheral blood leukocytes. Clinical immunology and immunopathology. 1986;38(2):150–60.

51. Garraud O, Borhis G, Badr G, Degrelle S, Pozzetto B, Cognasse F, et al. Revisiting the B-cell compartment in mouse and humans: more than one B-cell subset exists in the marginal zone and beyond. BMC immunology. 2012;13:63.

52. Caldwell HD, Kromhout J, Schachter J. Purification and partial characterization of the major outer membrane protein of Chlamydia trachomatis. Infection and immunity. 1981;31(3):1161–76.

53. Su H, Watkins NG, Zhang YX, Caldwell HD. Chlamydia trachomatis-host cell interactions: role of the chlamydial major outer membrane protein as an adhesin. Infection and immunity. 1990;58(4):1017–25.

54. Massari P, Toussi DN, Tifrea DF, de la Maza LM. Toll-like receptor 2-dependent activity of native major outer membrane protein proteosomes of Chlamydia trachomatis. Infection and immunity. 2013;81(1):303–10.

55. De Michele C, De Los Rios P, Foffi G, Piazza F. Simulation and Theory of Antibody Binding to Crowded Antigen-Covered Surfaces. PLoS computational biology. 2016;12(3):e1004752.

56. Tzeng SJ, Li WY, Wang HY. FcgammaRIIB mediates antigen-independent inhibition on human B lymphocytes through Btk and p38 MAPK. Journal of biomedical science. 2015;22:87.

57. Boeglin E, Smulski CR, Brun S, Milosevic S, Schneider P, Fournel S. Toll-like receptor agonists synergize with CD40L to induce either proliferation or plasma cell differentiation of mouse B cells. PloS one. 2011;6(10):e25542.

58. Rawlings DJ, Schwartz MA, Jackson SW, Meyer-Bahlburg A. Integration of B cell responses through Toll-like receptors and antigen receptors. Nature reviews Immunology. 2012;12(4):282–94.

59. Rosa D, Saletti G, De Gregorio E, Zorat F, Comar C, D’Oro U, et al. Activation of naive B lymphocytes via CD81, a pathogenetic mechanism for hepatitis C virus-associated B lymphocyte disorders. Proceedings of the National Academy of Sciences of the United States of America. 2005;102(51):18544–9.

60. Radaev S, Zou Z, Tolar P, Nguyen K, Nguyen A, Krueger PD, et al. Structural and functional studies of Igalphabeta and its assembly with the B cell antigen receptor. Structure (London, England : 1993). 2010;18(8):934–43.

61. Astsaturov IA, Matutes E, Morilla R, Seon BK, Mason DY, Farahat N, et al. Differential expression of B29 (CD79b) and mb-1 (CD79a) proteins in acute lymphoblastic leukaemia. Leukemia. 1996;10(5):769–73.

62. Hoogeboom R, Tolar P. Molecular Mechanisms of B Cell Antigen Gathering and Endocytosis. Current topics in microbiology and immunology. 2016;393:45–63.

63. Corneth OB, Klein Wolterink RG, Hendriks RW. BTK Signaling in B Cell Differentiation and Autoimmunity. Current topics in microbiology and immunology. 2016;393:67–105.

64. Okkenhaug K, Burger JA. PI3K Signaling in Normal B Cells and Chronic Lymphocytic Leukemia (CLL). Current topics in microbiology and immunology. 2016;393:123–42.

65. Bryan MA, Guyach SE, Norris KA. Specific humoral immunity versus polyclonal B cell activation in Trypanosoma cruzi infection of susceptible and resistant mice. PLoS neglected tropical diseases. 2010;4(7):e733.

66. Pone EJ, Zhang J, Mai T, White CA, Li G, Sakakura JK, et al. BCR-signalling synergizes with TLR-signalling for induction of AID and immunoglobulin class-switching through the non-canonical NF-kappaB pathway. Nature communications. 2012;3:767.

67. Donahue AC, Fruman DA. Proliferation and survival of activated B cells requires sustained antigen receptor engagement and phosphoinositide 3-kinase activation. Journal of immunology (Baltimore, Md : 1950). 2003;170(12):5851–60.

68. Bekeredjian-Ding I, Inamura S, Giese T, Moll H, Endres S, Sing A, et al. Staphylococcus aureus protein A triggers T cell-independent B cell proliferation by sensitizing B cells for TLR2 ligands. Journal of immunology (Baltimore, Md : 1950). 2007;178(5):2803–12.

69. Singh K, Bayrak B, Riesbeck K. A role for TLRs in Moraxella-superantigen induced polyclonal B cell activation. Frontiers in bioscience (Scholar edition). 2012;4:1031–43.

70. Pore D, Mahata N, Pal A, Chakrabarti MK. 34 kDa MOMP of Shigella flexneri promotes TLR2 mediated macrophage activation with the engagement of NF-kappaB and p38 MAP kinase signaling. Molecular immunology. 2010;47(9):1739–46.

71. Zhou H, Huang Q, Li Z, Wu Y, Xie X, Ma K, et al. PORF5 plasmid protein of Chlamydia trachomatis induces MAPK-mediated pro-inflammatory cytokines via TLR2 activation in THP-1 cells. Science China Life sciences. 2013;56(5):460–6.

72. Bas S, Neff L, Vuillet M, Spenato U, Seya T, Matsumoto M, et al. The proinflammatory cytokine response to Chlamydia trachomatis elementary bodies in human macrophages is partly mediated by a lipoprotein, the macrophage infectivity potentiator, through TLR2/TLR1/TLR6 and CD14. Journal of immunology (Baltimore, Md : 1950). 2008;180(2):1158–68.

73. Vaure C, Liu Y. A comparative review of toll-like receptor 4 expression and functionality in different animal species. Frontiers in immunology. 2014;5:316.

74. Ketloy C, Engering A, Srichairatanakul U, Limsalakpetch A, Yongvanitchit K, Pichyangkul S, et al. Expression and function of Toll-like receptors on dendritic cells and other antigen presenting cells from non-human primates. Veterinary immunology and immunopathology. 2008;125(1-2):18–30.

75. Karaki S, Anson M, Tran T, Giusti D, Blanc C, Oudard S, et al. Is There Still Room for Cancer Vaccines at the Era of Checkpoint Inhibitors. Vaccines. 2016;4(4).

76. Vinay DS, Kwon BS. Immunotherapy of cancer with 4-1BB. Molecular cancer therapeutics. 2012;11(5):1062–70.

77. Croft M. Control of immunity by the TNFR-related molecule OX40 (CD134). Annual review of immunology. 2010;28:57–78.

78. Vasir B, Zarwan C, Ahmad R, Crawford KD, Rajabi H, Matsuoka K, et al. Induction of antitumor immunity ex vivo using dendritic cells transduced with fowl pox vector expressing MUC1, CEA, and a triad of costimulatory molecules (rF-PANVAC). Journal of immunotherapy (Hagerstown, Md : 1997). 2012;35(7):555–69.

79. Robert C, Klein C, Cheng G, Kogan A, Mulligan RC, von Andrian UH, et al. Gene therapy to target dendritic cells from blood to lymph nodes. Gene therapy. 2003;10(17):1479–86.

80. RM J, H Y, NO S, K K, Y Z, RC B. B Cell Presentation of Chlamydia Antigen Selects Out Protective CD4γ13 T Cells: Implications for Genital Tract Tissue-Resident Memory Lymphocyte Clusters. Infection and immunity. 2018;86(2).

81. Cocco M, Stephenson S, Care MA, Newton D, Barnes NA, Davison A, et al. In vitro generation of long-lived human plasma cells. Journal of immunology (Baltimore, Md : 1950). 2012;189(12):5773–85.

82. E B, P K. Lymphocyte-polarized DC1s: Effective Inducers of Tumor-Specific CTLs. Oncoimmunology. 2012;1(8).

83. ML H, L H, C P, P I. Interdisciplinary Critique of sipuleucel-T as Immunotherapy in Castration-Resistant Prostate Cancer. Journal of the National Cancer Institute. 2012;104(4).

84. Carreno BM, Magrini V, Becker-Hapak M, Kaabinejadian S, Hundal J, Petti AA, et al. Cancer immunotherapy. A dendritic cell vaccine increases the breadth and diversity of melanoma neoantigen-specific T cells. Science (New York, NY). 2015;348(6236):803–8.

85. De Vries IJ, Krooshoop DJ, Scharenborg NM, Lesterhuis WJ, Diepstra JH, Van Muijen GN, et al. Effective migration of antigen-pulsed dendritic cells to lymph nodes in melanoma patients is determined by their maturation state. Cancer research. 2003;63(1):12–7.

86. Radomski M, Zeh HJ, Edington HD, Pingpank JF, Butterfield LH, Whiteside TL, et al. Prolonged intralymphatic delivery of dendritic cells through implantable lymphatic ports in patients with advanced cancer. Journal for immunotherapy of cancer. 2016;4:24.

87. Cheever MA, Higano CS. PROVENGE (Sipuleucel-T) in prostate cancer: the first FDA-approved therapeutic cancer vaccine. Clinical cancer research : an official journal of the American Association for Cancer Research. 2011;17(11):3520–6.

88. Wennhold K, Weber TM, Klein-Gonzalez N, Thelen M, Garcia-Marquez M, Chakupurakal G, et al. CD40-activated B cells induce anti-tumor immunity in vivo. Oncotarget. 2017;8(17):27740–53.

89. Lappin MB, Weiss JM, Delattre V, Mai B, Dittmar H, Maier C, et al. Analysis of mouse dendritic cell migration in vivo upon subcutaneous and intravenous injection. Immunology. 1999;98(2):181–8.

90. Creusot RJ, Yaghoubi SS, Chang P, Chia J, Contag CH, Gambhir SS, et al. Lymphoid-tissue-specific homing of bone-marrow-derived dendritic cells. Blood. 2009;113(26):6638–47.

91. Tanaka A, Sakaguchi S. Regulatory T cells in cancer immunotherapy. Cell research. 2017;27(1):109–18.

92. Ostrand-Rosenberg S, Fenselau C. Myeloid-Derived Suppressor Cells: Immune-Suppressive Cells That Impair Antitumor Immunity and Are Sculpted by Their Environment. Journal of immunology (Baltimore, Md : 1950). 2018;200(2):422–31.

