## Supplemental Material for "*Chlamydia*-induced *ex vivo* activation of B cells from mice, nonhuman primates, and humans produces large numbers of uniformly activated antigen presenting cells"

**Figure S1. Generation of human CAB and effect of prior _Chlamydia_ infection on CAB generation.** A) Fluorescently labeled PBMCs were incubated with inactivated *Ct* $L_2$ EB for 4 days or left untreated, and B cell proliferation analyzed by flow cytometry. Top panels show representative contour plots delineating the gating strategies, while lower panels show characterization of human resting B cells (blue) and CABs (red). B) Dose-response curves of the proliferative response of human B cells to stimulation with inactivated *Ct* serovar D EB and live *Ct* serovar $L_2$ EB (mean ±SD). C) Dose-response curves of proliferative response to stimulation with inactivated *C. trachomatis* serovar $L_2$ EB of human B cells from donors that did not display (termed uninfected) or did display *Chlamydia*-specific T cell responses (termed *Ct* infected) (mean ± SD). D) Representative contour plots of human resting B cells and CAB showing CD19/CD20 expression levels (left panel) and quantitative comparison of their respective MFI (right panel). E) Prior genital *Chlamydia* infection did not affect generation of NHP CAB. RM PBMC were isolated and cryopreserved before and after animals had been genitally inoculated with *Ct* serovar D. PBMC were thawed, fluorescently labeled, and incubated with varying amounts of inactivated *Ct* $L_2$ EB. Dose-response curves of the *Chlamydia*-induced proliferative response of NHP B cells before and after *Ct* infection are shown (mean ±SD). F) *Chlamydia* infection status did not significantly affect ability of CAB to prime antigen-specific $CD8^+$ T cell responses *in vivo* in mice. $10^6$ fluorescently labeled OT-I cells were IV transferred into B6 mice that had been genitally infected with *Ct* serovar $L_2$ 2 months prior or remained uninfected. The following day, $10^7$ unloaded or OVA-loaded CAB were IV transferred. Mice were euthanized 3 days later, and spleens excised and processed into single-cell suspensions to evaluate degree of OT-I proliferation by flow cytometry. Left panels show representative heat map statistic plots (color scale indicates level of CD44), right panel display quantification of total OT-I cells/spleen (mean ±SD). G) Human CAB do not harbor infectious *Chlamydia.* IFUs measured in the stock of *C. trachomatis* serovar $L_2$ EB used in (B) to generate human CAB (top panel) and in CAB generated with this stock, which were lysed by sonication immediately before IFU determination (lower panel).

**Figure S2. Generating murine or NHP CAB and identification of human TLR activated by inactivated *Ct* $L_2$ EB.** A) Fluorescently labeled murine splenocytes from naïve B6 mice were incubated with inactivated *Ct* $L_2$ EB for 4 days. Representative contour plots depict gating strategy used to assess B cell proliferation by flow cytometry. B) Representative contour plots characterize murine resting B cells (blue) and CAB (red). C) Dose-response curves of the proliferative response of murine B cells to stimulation with inactivated EB from *Ct* serovar D or *C. muridarum* (mean ±SD). D) Representative contour plots and bar graphs illustrate increased expression of CD45R and MHC-II by murine CAB (red) compared to murine resting B cells (blue). E) Representative heat map statistic plots reveals the limited number of T cells, myeloid cells, and NK cells in murine CAB cultures (color scale indicates level of SSC-A). F) Generation of NHP CAB. Fluorescently labeled PBMC from RM were cultured with inactivated *Ct* $L_2$ EB. Representative contour plots depict gating strategy used to define RM resting B cells (blue) and CAB (red). G) TLR screening was performed using a panel of HEK293-TLR-Blue clones engineered to express only a single specific TLR and a SEAP–reporter plasmid activated with NF-κB, AP-1, and IRF7 transcription factors. Cells incubated with indicated TLR-specific ligands were used as positive controls. Cell activation was defined as an increase in SEAP activity measured as absorbance (O.D.) at 650 nm, using QUANTI-Blue reagent, according to the manufacturer's protocol.

**Figure S3. Non-antigen specific BCR signaling is involved in generating CAB that retain function after cryopreservation.** A) Heat inactivation or proteolysis eliminated ability of *Ct* to induce proliferation of murine and human B cells. Inactivated *Ct* $L_2$ EB incubated for 30 minutes at 56°C, treated with proteinase K, or left untreated were used to stimulate murine splenocytes (A) or human PBMC (B) as indicated in Figure 1. Dose-response curves for both species are shown. B) BCR crosslinking or TLR2 signaling alone did not induce proliferation of human B cells. Fluorescently labeled PBMC were incubated with inactivated *Ct* $L_2$ EB (*Ct*), anti-human IgM+IgG+IgA F(ab')$_2$ fragments ($\alpha$-BCR), or Pam$_3$CSK$_4$ (TLR2/1 agonist) for 4 days or left untreated. B cell proliferation was defined by flow cytometry (representative contour plots are shown). C) Inhibition of Syk-dependent, but not MAPK-dependent, signaling blocked *Ct*-induced activation of human B cells. Fluorescently-labeled human PBMC were pre-incubated with BIIB-057 (Syk inhibitor) or PD98059 (MAPK kinase inhibitor) for 30 minutes or left untreated before exposure to inactivated *Ct* $L_2$ EB for 4 days. B cell proliferation was analyzed by flow cytometry (normalized proliferation shown). Data displayed from 3 independent experiments shown, bars indicate mean $\pm$SD. D) Cryopreservation did not alter CAB purity or activation marker expression. CAB from B6 mice that were freshly generated or previously cryopreserved as indicated in Methods were stained for flow cytometric evaluation. Gating strategy used to evaluate CAB, showing representative contour plots from cryopreserved CAB. (E) Representative contour plots depict CD45R and MHC-II expression by freshly generated or previously cryopreserved murine CAB (C57BL/6J), demonstrating comparable between-group frequency of CD45R and MHC-II expression and near complete absence of MHC-II positive cells. (F) Representative contour plots depict expression of CD40, CD54, CD86 and MHC-II by freshly generated and previously cryopreserved CAB generated using splenocytes from 2 commonly used mouse strains (C57BL/6J and BALB/cJ).

**Figure S4. Phenotypic characterization of human and NHP CAB and gating strategies used to define splenic FoxP3$^+$ regulatory T cells and MDSC in tumor-bearing mice.** A) Human PBMC were stimulated with *Ct* as indicated in Figure 1. Representative contour plots and histograms compare CD2 expression in resting B cells, B cells activated with *Ct* (CAB) or CpG, and T cells. B) Bar graph shows CD2 iMFI levels for cell populations depicted in (A). C) Representative contour plots and histograms comparing the expression of CD14, CD16 and CD66abce by B cells treated as indicated in (A), and monocytes from PBMC stimulated with CpG for 4 days. D) Bar graph shows CD66abce iMFI levels of cell populations depicted in (C). E) NHP PBMC were stimulated with *Ct* or left untreated as indicated in Figure 3. Representative contour plots and histograms comparing the expression of CD2 in resting B cells, CAB, and T cells. F) Bar graph shows CD2 iMFI levels of cell populations depicted in (E). G) Gating strategy used to evaluate NHP PBMC stimulated with *Ct*. H) Representative contour plots depict purity of NHP CAB that routinely contained >98% CAB after negative immunomagnetic (the majority of contaminating cells were HLA-DR negative). I-J) Representative contour plots of gating strategy used to define splenic FoxP3$^+$ regulatory T cells (I) and MDSC (J) in tumor-bearing mice administered unloaded CAB or CAB loaded with tumor-associated antigen. Representative heatmap statistic plots show distribution of Gr-1 (color scale) in myeloid cells and polymorphonuclear and monocytic MDSC.

**Figure S5. Western blots assessing phosphorylation of Akt and Btk in human B cells exposed to *Ct*.** Human B cells (purified from PBMC by negative immunomagnetic selection), were stimulated with *Ct* or vehicle for 15 minutes and western blot assessed the phosphorylation of Akt (A) and Btk (B). Unprocessed original images of western blots for detection of total Akt and Btk as well as for detection of phosphorylation of Btk and Akt are shown. Inh, indicates use of LY294002, a potent inhibitor of the PI3K/AKT pathway.

### Figure S1

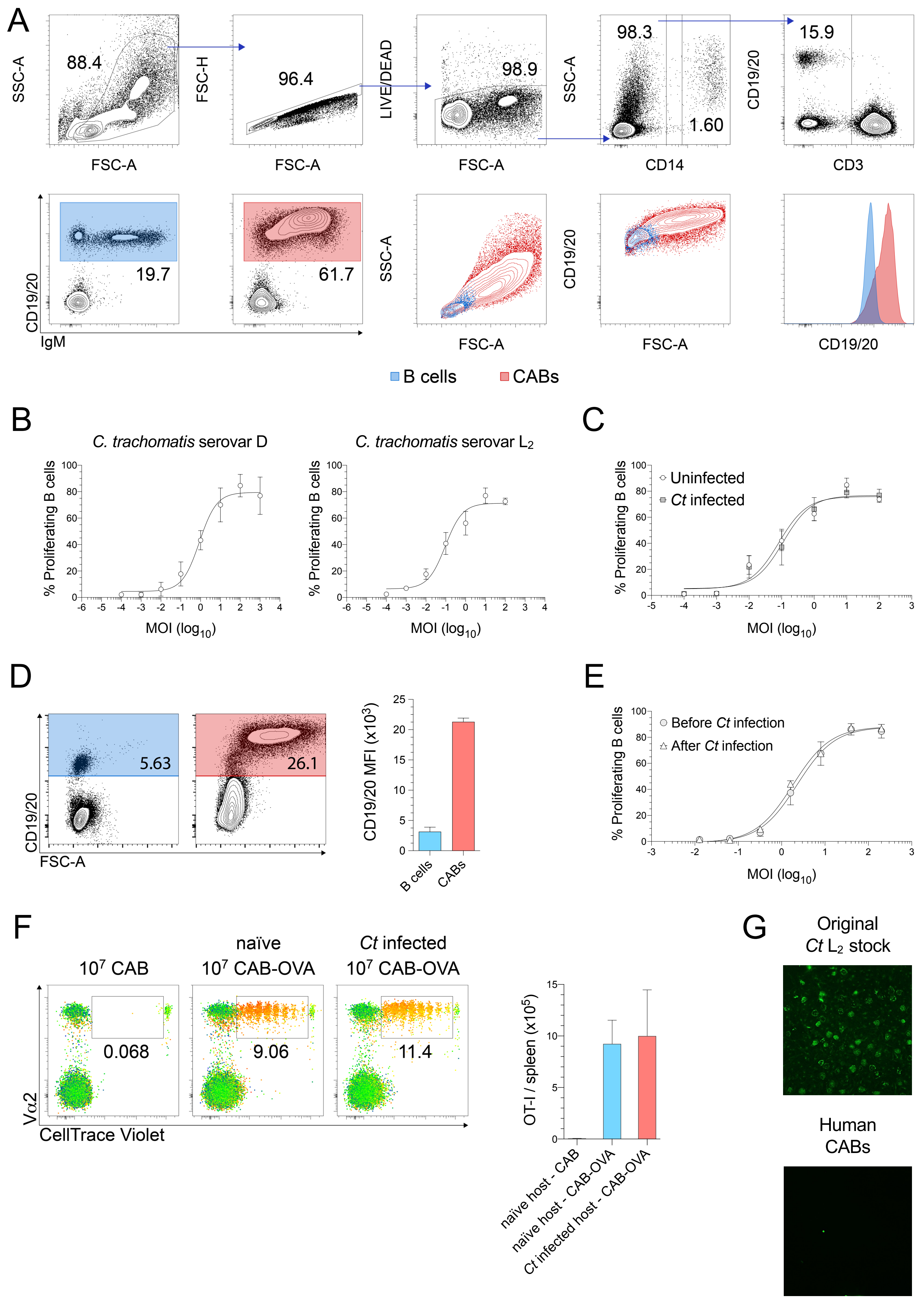
A
SSC-A
FSC-A
88.4
FSC-H
96.4
LIVE/DEAD
98.9
98.3
1.60
CD14
CD19/20
15.9
CD3
CD19/20
IgM
19.7
61.7
B cells
CABs
B
C. trachomatis serovar D
C. trachomatis serovar L2
% Proliferating B cells
MOI (log10)
C
Uninfected
Ct infected
D
5.63
26.1
FSC-A
CD19/20 MFI (x10^3)
E
Before Ct infection
After Ct infection
F
10^7 CAB
naïve 10^7 CAB-OVA
Ct infected 10^7 CAB-OVA
0.068
9.06
11.4
Vα2
CellTrace Violet
OT-I / spleen (x10^5)
naïve host - CAB
naïve host - CAB-OVA
Ct infected host - CAB-OVA
G
Original Ct L2 stock
Human CABs

### Figure S2

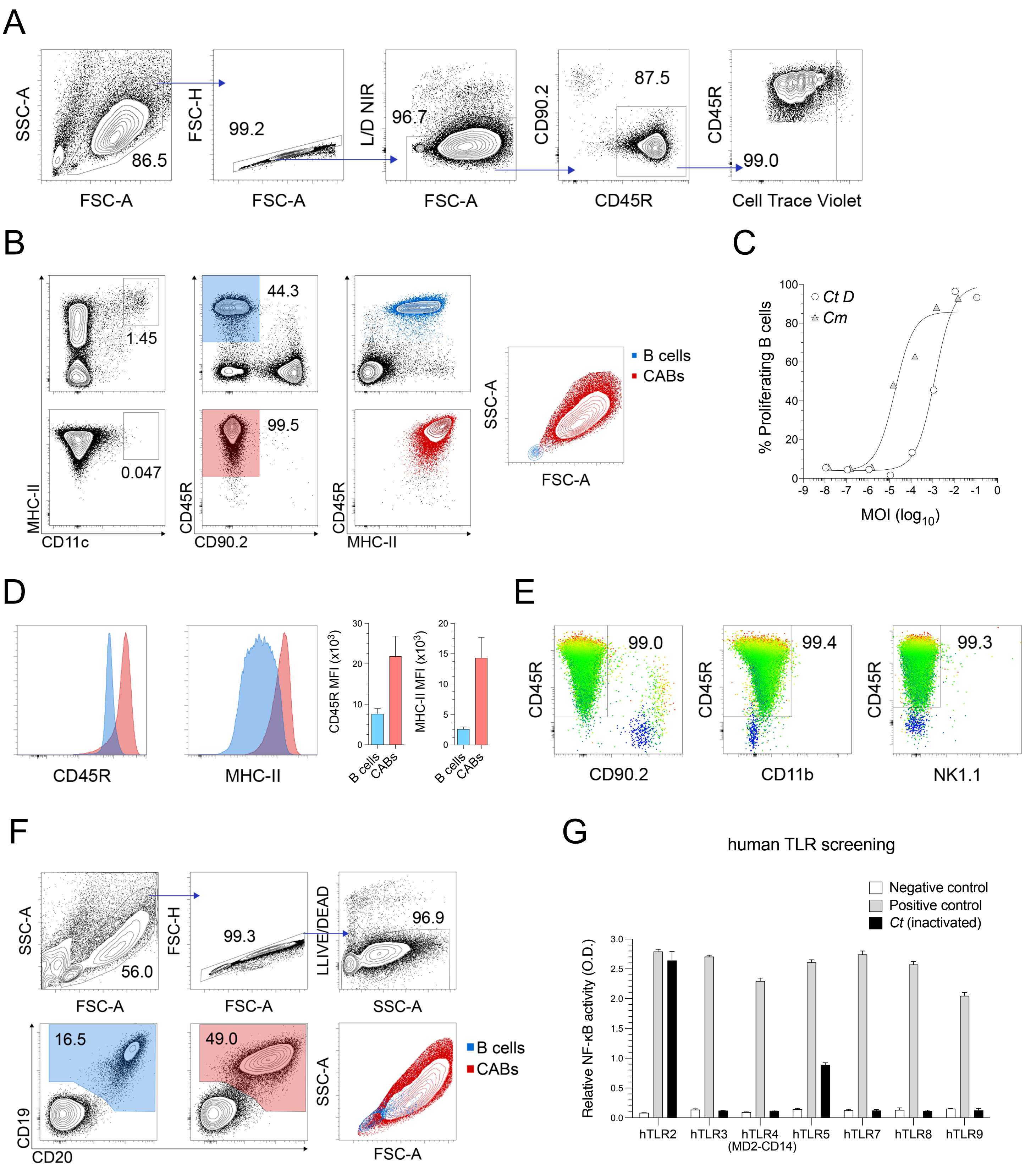


Positive controls

hTLR2: HKLM (heat-killed Listeria monocytogenes, $1x10^8$ cells/mL)
hTLR3: Poly(I:C) HMW (1 µg/mL)
hTLR4: E. coli K12 LPS (100 ng/mL)
hTLR5: S. typhimurium flagellin (100 ng/mL)
hTLR7: CL097 (1 µg/mL)
hTLR8: CL075 (1 µg/mL)
hTLR9: CpG ODN2006 (1 µg/mL)

### Figure S3

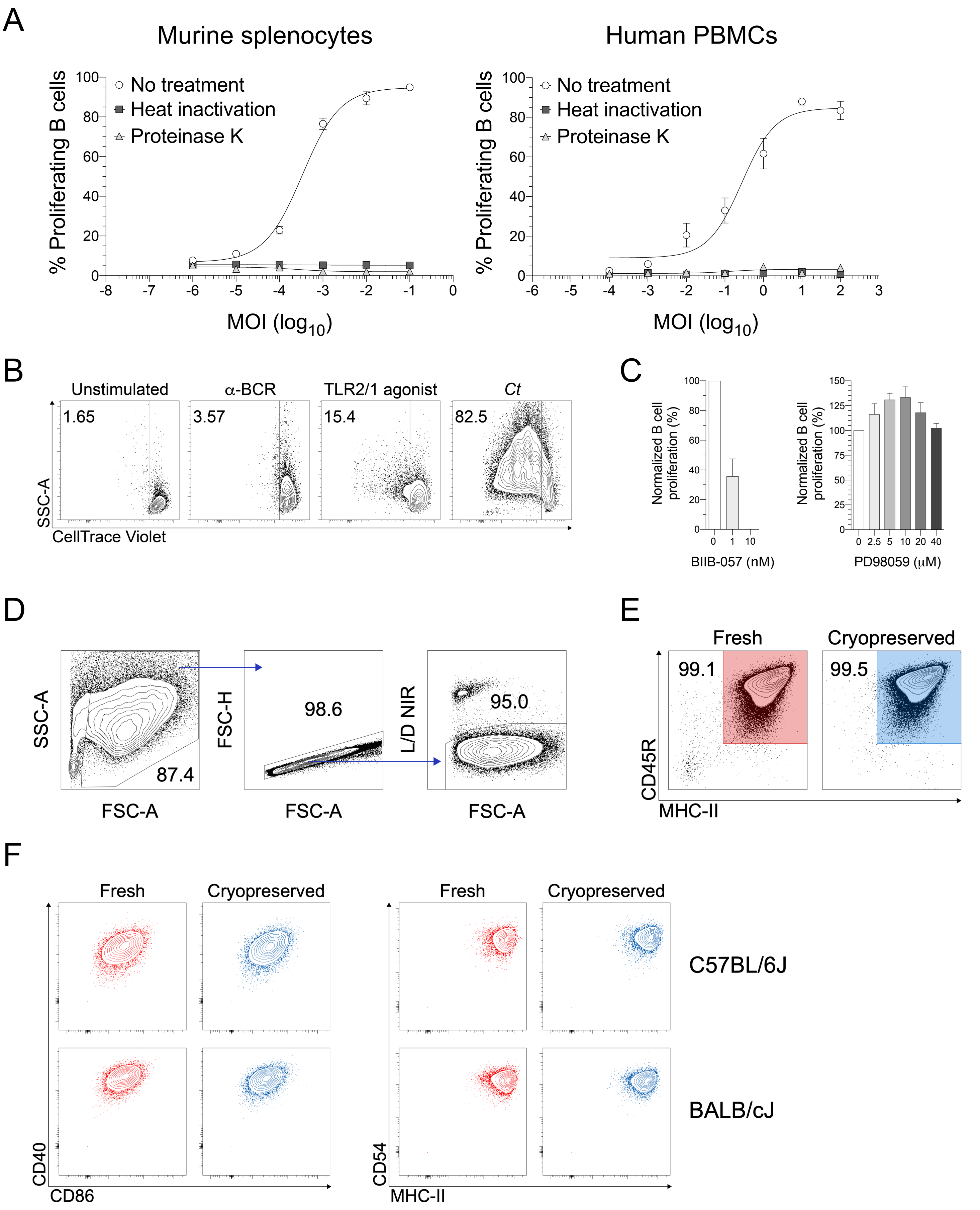
A
Murine splenocytes
Human PBMCs
% Proliferating B cells
No treatment
Heat inactivation
Proteinase K
MOI (log10)
B
Unstimulated
α-BCR
TLR2/1 agonist
Ct
1.65
3.57
15.4
82.5
SSC-A
CellTrace Violet
C
Normalized B cell proliferation (%)
BIIB-057 (nM)
PD98059 (μM)
D
SSC-A
FSC-A
87.4
FSC-H
98.6
L/D NIR
95.0
E
Fresh
Cryopreserved
99.1
99.5
CD45R
MHC-II
F
Fresh
Cryopreserved
C57BL/6J
BALB/cJ
CD40
CD86
CD54
MHC-II

### Figure S4

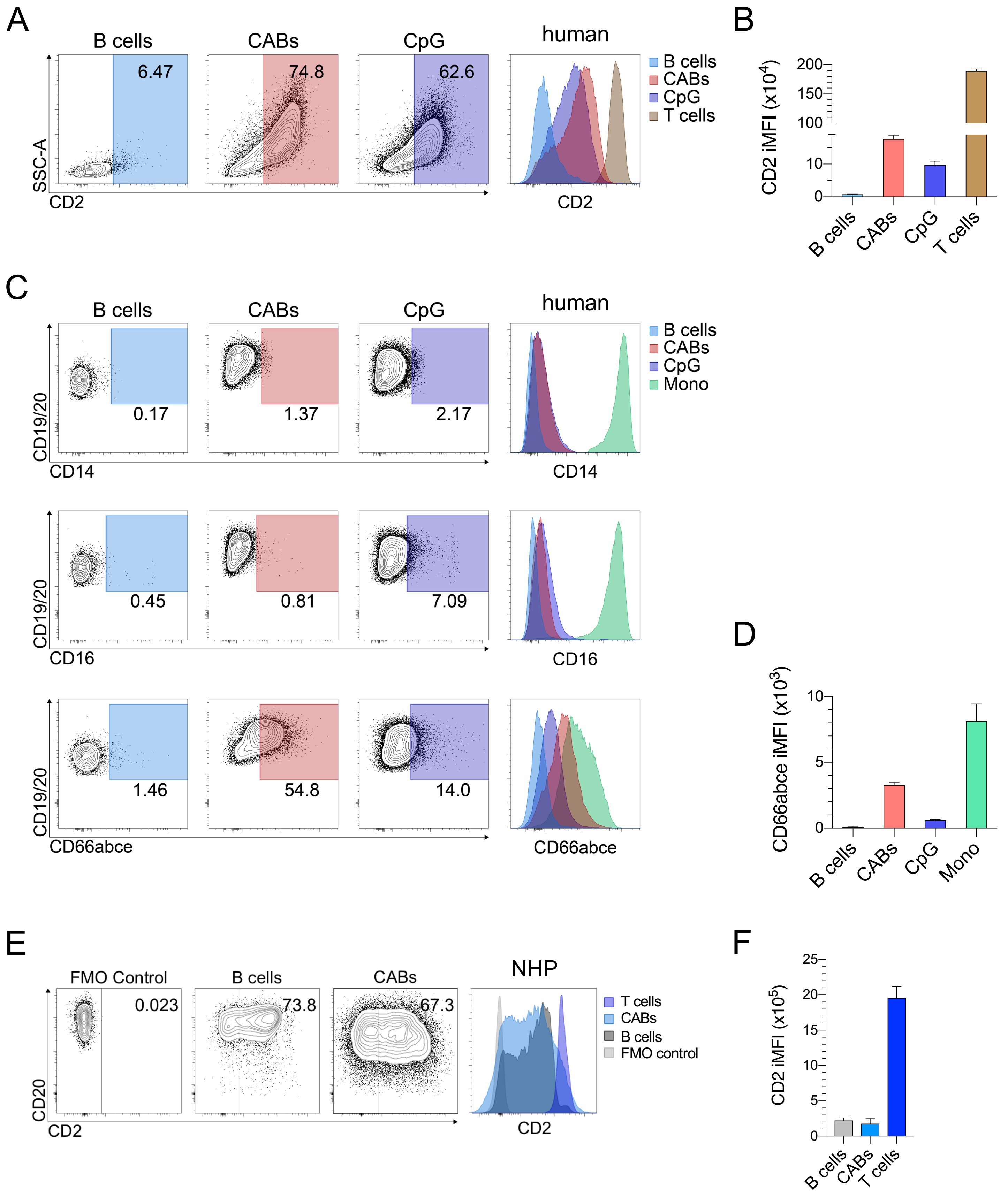
A
B cells
CABs
CpG
human
6.47
74.8
62.6
SSC-A
CD2
B cells
CABs
CpG
T cells
CD2
B
CD2 iMFI (x10$^4$)
200
150
100
10
0
B cells
CABs
CpG
T cells
C
B cells
CABs
CpG
human
0.17
1.37
2.17
CD19/20
CD14
B cells
CABs
CpG
Mono
CD14
0.45
0.81
7.09
CD19/20
CD16
CD16
1.46
54.8
14.0
CD19/20
CD66abce
CD66abce
D
CD66abce iMFI (x10$^3$)
10
5
0
B cells
CABs
CpG
Mono
E
FMO Control
B cells
CABs
NHP
0.023
73.8
67.3
CD20
CD2
T cells
CABs
B cells
FMO control
CD2
F
CD2 iMFI (x10$^5$)
25
20
15
10
5
0
B cells
CABs
T cells

### Figure S4

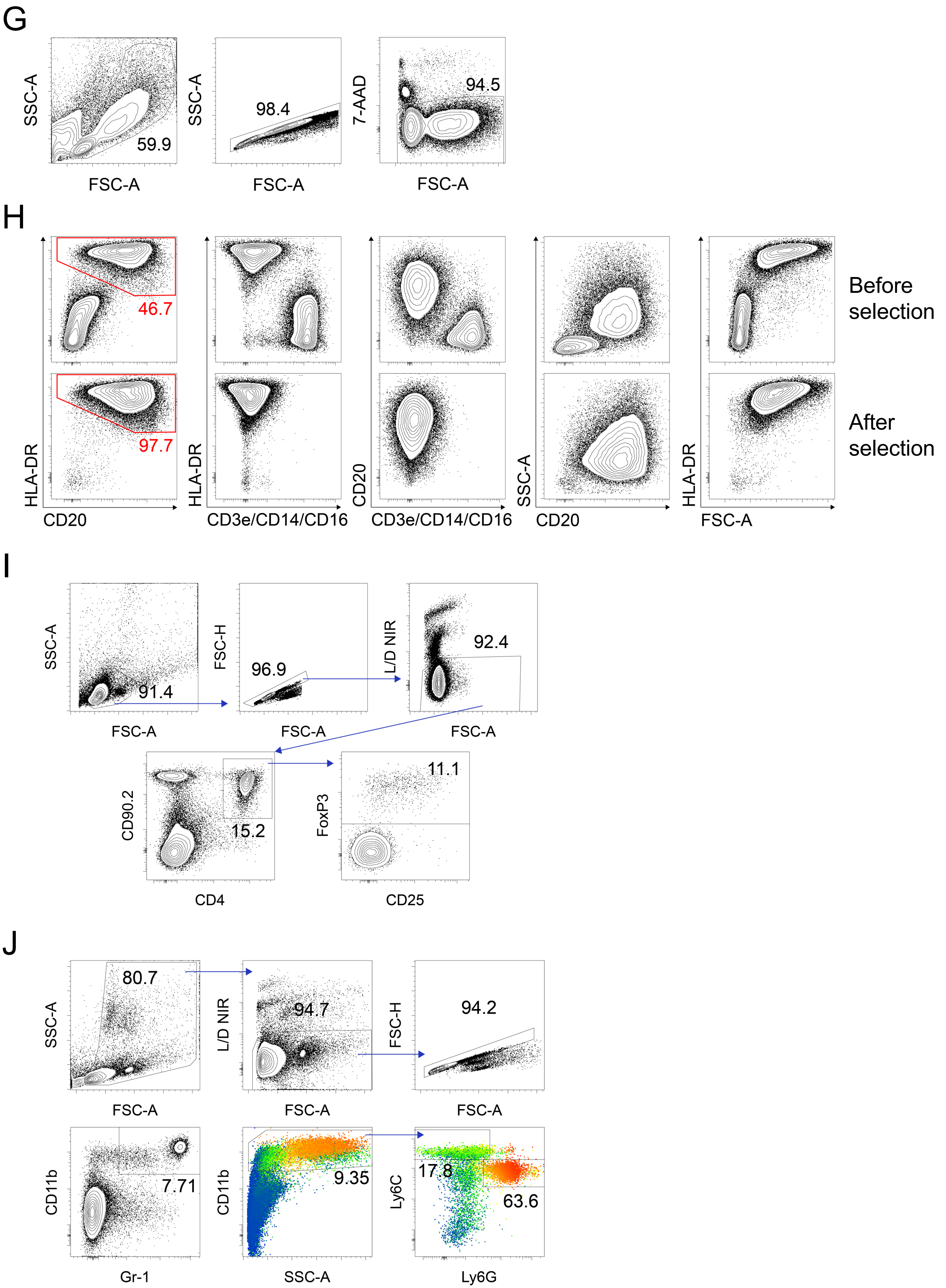
G
SSC-A
FSC-A
59.9
98.4
7-AAD
94.5
H
HLA-DR
CD20
46.7
97.7
CD3e/CD14/CD16
SSC-A
FSC-A
Before selection
After selection
I
SSC-A
FSC-A
91.4
FSC-H
96.9
L/D NIR
92.4
CD90.2
CD4
15.2
FoxP3
CD25
11.1
J
SSC-A
FSC-A
80.7
L/D NIR
94.7
FSC-H
94.2
CD11b
Gr-1
7.71
SSC-A
9.35
Ly6C
Ly6G
17.8
63.6

### Figure S5

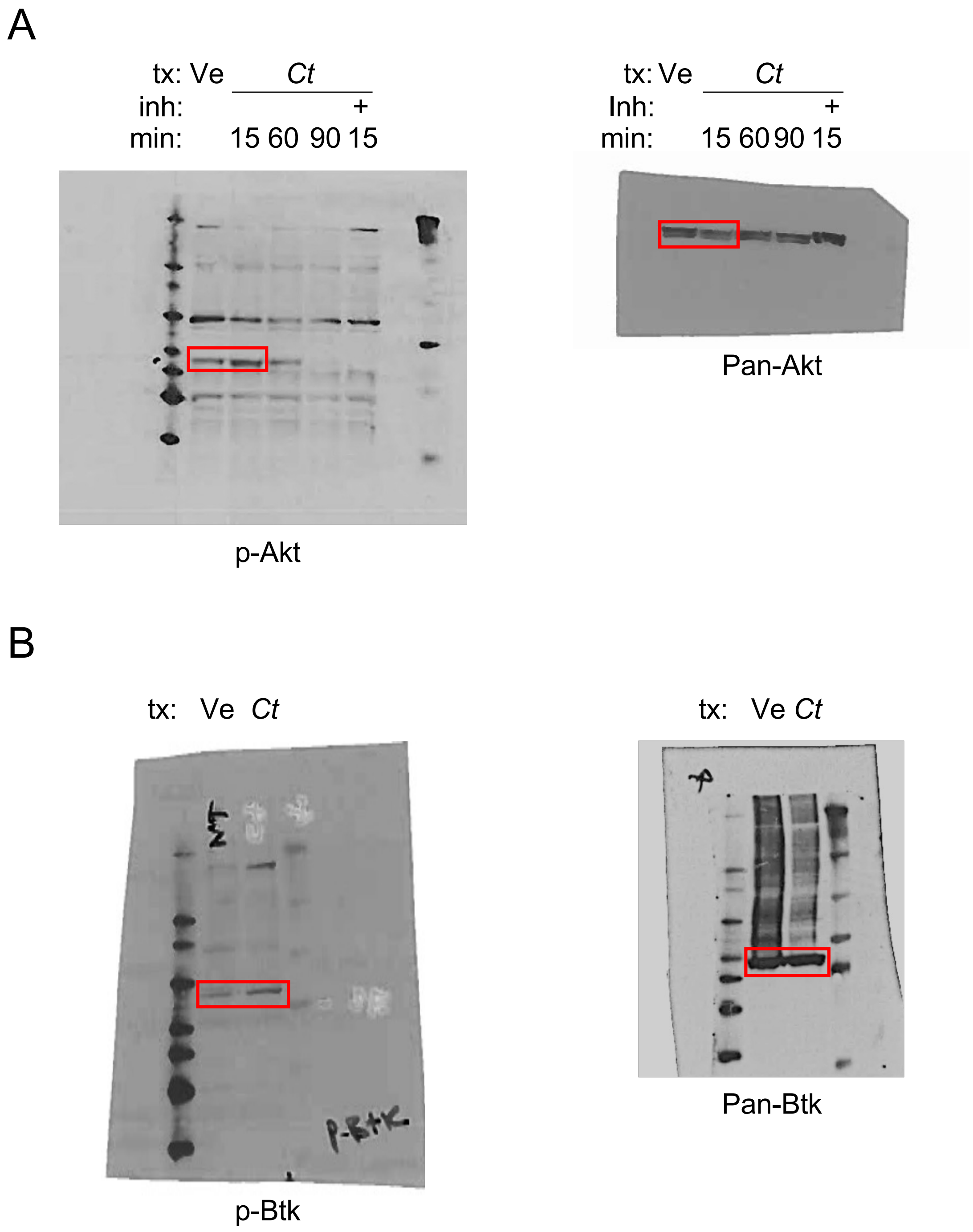

A
tx: Ve Ct
inh: +
min: 15 60 90 15
p-Akt
tx: Ve Ct
Inh: +
min: 15 60 90 15
Pan-Akt
B
tx: Ve Ct
p-Btk
tx: Ve Ct
Pan-Btk
